# Contiguous and complete assemblies of *Blastocystis* gut microbiome-associated protists reveal evolutionary diversification to host ecology

**DOI:** 10.1101/2023.11.20.567959

**Authors:** Abigail L. Lind, Nathan A. McDonald, Elias R. Gerrick, Ami S. Bhatt, Katherine S. Pollard

**Affiliations:** Gladstone Institute for Data Science and Biotechnology, Gladstone Institutes, San Francisco, CA; School of Biological Sciences, Georgia Institute of Technology, Atlanta, Georgia; Department of Biology, Stanford University, Stanford, CA; Department of Microbiology, University of Chicago, Chicago, IL; Department of Genetics, Stanford University, Stanford, California, USA; Department of Medicine (Hematology, Blood and Marrow Transplantation), Stanford University, Stanford, California, USA; Department of Epidemiology & Biostatistics, University of California, San Francisco, CA; Chan Zuckerberg Biohub SF, San Francisco, CA

**Author notes:** **Corresponding author contact information** 1700 Owens Street, San Francisco, CA 94158.

## Abstract

*Blastocystis*, an obligate host-associated protist, is the most common microbial eukaryote in the human gut and is widely distributed across vertebrate hosts. The evolutionary transition of *Blastocystis* from its free-living stramenopile ancestors to a radiation of host-associated organisms is poorly understood. To explore this, we cultured and sequenced eight strains representing the significant phylogenetic diversity of the genus using long-read, short-read, and Hi-C DNA sequencing, alongside gene annotation and RNA sequencing. Comparative genomic analyses revealed significant variation in gene content and genome structure across *Blastocystis.* Notably, three strains from herbivorous tortoises, phylogenetically distant from human subtypes, have markedly larger genomes with longer introns and intergenic regions, and retain canonical stop codons absent in the human-associated strains. Despite these genetic differences, all eight isolates exhibit gene losses linked to the reduced cellular complexity of *Blastocystis,* including losses of cilia and flagella genes, microtubule motor genes, and signal transduction genes. Isolates from herbivorous tortoises contained higher numbers of plant carbohydrate-metabolizing enzymes, suggesting that like gut bacteria, these protists ferment plant material in the host gut. We find evidence that some of these carbohydrate-metabolizing enzymes were horizontally acquired from bacteria, indicating that horizontal gene transfer is an ongoing process in *Blastocystis* that has contributed to host-related adaptation. Together, these results highlight substantial genetic and metabolic diversity within the *Blastocystis* genus, indicating different lineages of *Blastocystis* have varied ecological roles in the host gut.

## Introduction

The vertebrate gut is host to a diverse ecosystem of bacteria, archaea, viruses, and microbial eukaryotes. This diverse microbial community has broad impacts on the physiology and development of the host. Gut microbes can metabolize compounds that are not digestible by the host, providing access to previously inaccessible nutrients (Yadav et al. 2018). Gut microbes also influence the host’s immune system and impact host health through their metabolic byproducts (Sommer and Bäckhed 2013). The microbiome’s composition, which is influenced by diet, physiology, and the host immune system, varies substantially across vertebrate animals (David et al. 2014; Thorman et al. 2023; Godoy-Vitorino et al. 2012).

Compared to bacteria, substantially less is understood about microbial eukaryote impacts on host physiology and the gut ecosystem, despite the prevalence and diversity of gut microbial eukaryotes across vertebrates (Campo et al. 2020; Nash et al. 2017). The protist *Blastocystis* is the most prevalent eukaryotic microorganism in the human gut, found in 10-50% of industrialized populations and at higher prevalence in non-industrialized communities (Khaled et al. 2020; Scanlan and Marchesi 2008; Scanlan et al. 2014; Asnicar et al. 2021; Jinatham et al. 2021; Scanlan and Stensvold 2013). Beyond humans, *Blastocystis* colonizes a wide range of animals including other mammals, birds, reptiles, fish, amphibians, and insects (Gantois et al. 2020; Boreham and Stenzel 1993; Yoshikawa et al. 2016). *Blastocystis* is a genetically diverse genus, with strains adapting to varied gut environments across different host species (Stensvold et al. 2007). Its presence is associated with positive health outcomes in humans, including lower levels of gut inflammation and metabolic health (Piperni et al. 2024; Asnicar et al. 2021; Nieves-Ramírez et al. 2018). However, very little is understood about how *Blastocystis* contributes to the gut environment and interacts with other gut microbiota.

*Blastocystis* belongs to the Opalinata lineage of stramenopiles, a group of protists that are obligate animal gut commensals (Kostka 2017). Unlike most stramenopiles, *Blastocystis* is non-motile, lacks flagella or surface hairs, and grows predominantly as a round cell with large central vacuole (Zierdt 1988). The *Blastocystis* genus contains numerous genetically distinct groups categorized into subtypes based on polymorphism in the 18S rRNA gene. Thus far, 40 distinct subtypes have been proposed, and this number is continuing to expand (Santin et al. 2024; Stensvold and Clark 2020). To date, strains from three axenic lab-cultured subtypes (ST1, ST4, and ST7), all of which are subtypes observed in humans, have been sequenced, annotated, and deposited in public databases (Denoeud et al. 2011; Gentekaki et al. 2017). While these subtypes span a narrow range of the *Blastocystis* phylogenetic diversity, they have very different genomes and gene contents (Gentekaki et al. 2017). These genomes have either gapped scaffolds (ST7) or highly fragmented non-contiguous genome assemblies (ST1, ST4), limiting the ability to determine synteny between subtypes, and two genomes were annotated without consideration for the unique *Blastocystis* phenomenon where a portion of genes lack DNA-encoded termination codons. Thus, we lack highly complete and well annotated genomes necessary for establishing cases of gene loss, horizontal transfer, and sequence evolution so that functional repertoires can be associated with strain diversification.

Like most host-associated eukaryotes, *Blastocystis* grows best in the presence of other microbiota. Establishing robust axenic cultures of intestinal protists including *Blastocystis* is laborious and often either fails to remove all microbial contaminants or results in the death of the protist (Zierdt and Williams 1974; Tan 2008). However, removing sources of microbial contamination is important for eukaryotic genome assembly via short-read sequencing (Gough et al. 2020). As a result, genome sequencing for *Blastocystis* and other intestinal protists has been limited to axenic cultures (Keeling et al. 2014; Gough et al. 2020). To overcome this barrier, we establish high quality and contamination-free genomes from xenic cultures of *Blastocystis* grown together with other microbes by using long-read Nanopore and Hi-C sequencing technologies coupled with short reads and metagenomic assembly techniques to establish genomes of *Blastocystis* strains that are grown together with other microbes. The longer read length of Nanopore sequencing helps span repetitive regions, resulting in greater contig length and fewer misassembles, and has been successfully applied to assembling bacterial metagenomes (Moss et al. 2020). Hi-C sequencing, which captures spatial interactions between DNA fragments, can further improve genome assembly by correcting misassemblies and scaffolding contigs (Šimková et al. 2024). By combining these technologies with metagenomic assembly techniques, we were able to generate nearly complete *Blastocystis* genomes without the need for axenic culturing. This approach allows us to investigate the full genetic diversity of *Blastocystis*, even in the presence of other microbes, and to assemble highly contiguous genomes that were previously difficult to obtain.

Here, we applied this hybrid sequencing strategy to eight *Blastocystis* strains, five isolated from humans and three isolated from the herbivorous tortoise *Testudo horsfeldii.* These isolates, five of which are xenic and three of which are axenic, span the phylogenetic diversity of the *Blastocystis* genus, enabling us to investigate *Blastocystis* evolution and determine how *Blastocystis* has adapted to thrive in animal hosts with different diets, physiologies, and resident microbiota. Our highly contiguous and complete genomes allowed us to investigate patterns of genome evolution within and between subtypes, and to compare the genus to related protists. This work demonstrates how high-quality genome assemblies can be created from xenic protist culture by combining deep long-read sequencing with metagenomic assembly techniques, enabling rigorous comparative genomic analyses.

## Results

### Capturing Blastocystis diversity

To investigate the evolution of the *Blastocystis* genus, from its initial divergence through its later diversification, we sequenced eight diverse *Blastocystis* strains. Strains were cultured anaerobically on egg media overlayed with a balanced salt solution and 25% heat-inactivated horse serum. Five of these strains correspond to 3 different previously defined “subtypes” (genetically similar groups historically identified in mammalian and avian species (Stensvold and Clark 2020). These five strains were originally cultured from human fecal samples and were obtained from the American Type Culture Collection. These human isolated strains correspond to three of the four most prevalent *Blastocystis* “subtypes” (Beghini et al. 2017), genetically similar groups historically identified in mammalian and avian species (Stensvold and Clark 2020): Two subtype (ST) 1 (Nandll, JDR), two ST3 (DL, NMH), and one ST4 (BT1). The remaining three strains were isolated from companion animal tortoises: two from different individuals of the domestic Russian tortoise species *Testudo horsfeldii* (Rus-B, Rus-S), and one from a Hermann’s tortoise *Testudo hermanni* (Hermann’s). We performed comparisons of the 18S rRNA gene of these our strains to other *Blastocystis* strains, finding that the tortoise isolates were distantly related to the human isolates and do not belong to a described mammalian or avian subtype (Figure 1A, Table S1). Analysis of all eight cultures by light microscopy revealed predominantly vacuolar *Blastocystis* cells, containing a large central vacuole and a thin peripheral ring of cytoplasm. While *Blastocystis* have been observed to adopt a variety of morphologies, vacuolar morphologies dominate cultured samples (Figure 1B).

**Figure 1.**
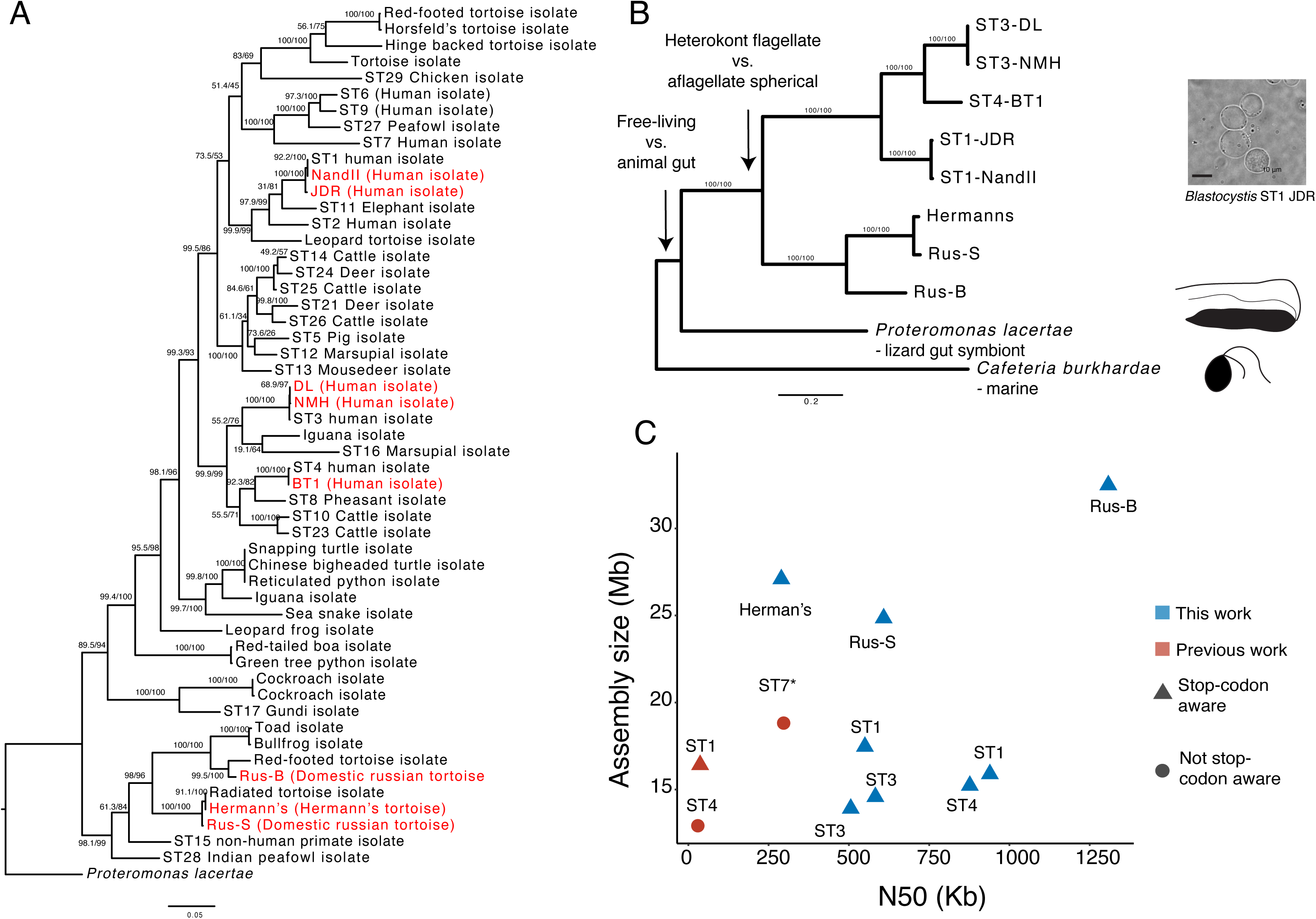
(A) 18S rRNA maximum likelihood (ML) phylogeny of Blastocystis subtypes, with *Proteromonas lacertae* as an outgroup. Support values are 1000 replicates of ultrafast bootstrap and SH-aLRT tests (Guindon et al. 2010; Hoang et al. 2018). Strains in red were sequenced in this study. (B) 13-gene concatenation ML phylogeny for all strains and species with sequenced genomes used in this study. Important evolutionary transitions are highlighted. Images represent cellular morphology of *Cafeteria burkhardae*, *Proteromonas lacertae*, and *Blastocystis*. (C) Assembly statistics of Blastocystis strains sequenced in this work compared with previously sequenced and annotated genomes. Stop-codon aware and not stop codon aware designations refer to gene annotation methodologies that take into consideration that Blastocystis genes can lack stop codons or do not consider this, respectively. The ST7 N50 is a gapped scaffold N50 generated with Sanger sequencing scaffolding.

To explore how the *Blastocystis* genus has emerged and diversified, we compared the *Blastocystis* genomes generated in this work to those of two other sequenced stramenopiles: *Cafeteria burkhardae*, a free-living heterotrophic marine flagellate (Hackl et al. 2020), and the recently sequenced *Proteromonas lacertae*, a reptile gut commensal closely related to *Blastocystis* (Záhonová et al. 2023). These organisms are the closest relatives to *Blastocystis* that have sequenced genomes. Genome comparisons allowed us to reconstruct three evolutionary transitions that have occurred during stramenopile evolution: the transition from a free-living aquatic lifestyle to an anaerobic lifestyle inside the animal gut (comparing *Cafeteria* to *Blastocystis*/*Proteromonas*), the change in cell morphology from heterokont flagellate to aflagellate, spherical cells (comparing *Cafeteria*/*Proteromonas* to *Blastocystis*), and adaptations to different animal hosts (comparing *Blastocystis* tortoise to *Blastocystis* human) (Figure 1B).

### Contiguous and complete Blastocystis genomes

We prepared high quality, high molecular weight DNA, as well as RNA, from the eight *Blastocystis* cultures. DNA was sequenced with Nanopore long-read technology. Short-read DNA and RNA sequencing was also performed for assembly polishing and gene annotation. Hi-C data was additionally generated to help resolve scaffolds for one strain (ST4-BT1). Non-eukaryotic sequence was detected via multiple methods and excluded from final assemblies (see Methods, Genome assembly and decontamination). We additionally validated that our genomes were not contaminated by bacterial sequence by determining that genes with high sequence similarity to bacterial genes arose from polyadenylated eukaryotic transcripts (Table S2) (see Methods, Gene contamination and horizontal gene transfer screens).

This combination of sequencing technologies enabled us to generate highly contiguous genomes for all six subtypes. The N50 of these assemblies ranged from 289 Kb to 1.3 Mb, achieving a much higher assembly quality than currently available *Blastocystis* genome datasets in NCBI (median: 37 Kb) (Table 1, Figure 1C). We used the BUSCO pipeline to estimate genome completeness based on presence and absence of single-copy universal eukaryotic genes (Manni et al. 2021). BUSCO completeness estimates ranged from 42% to 61.2%, similar to BUSCO completeness results of previously sequenced *Blastocystis* isolates and related species (Table 1, Figure 1C). These values likely underestimate the true completeness, because BUSCO is known to have poorer performance in protists relative to plants, animals, and fungi (Saary et al. 2020). Nonetheless, the consistent recovery of a similar fraction of BUSCO genes across stramenopile genomes indicates comparable completeness. It is therefore striking that the sequenced *Blastocystis* strains showed substantial variability in genome size, from 14.0 to 32.2 Mb. Strains in the same subtype exhibited similar lengths: ST3 isolates were 14.0 and 15.3 Mb, and ST1 isolates were 17.2 and 16.0 Mb in size. In contrast, the genomes of the three tortoise isolates were ∼1.5 to 2 times larger than human isolates, ranging from 26.0 Mb to 33.3 Mb.

**Table 1.**
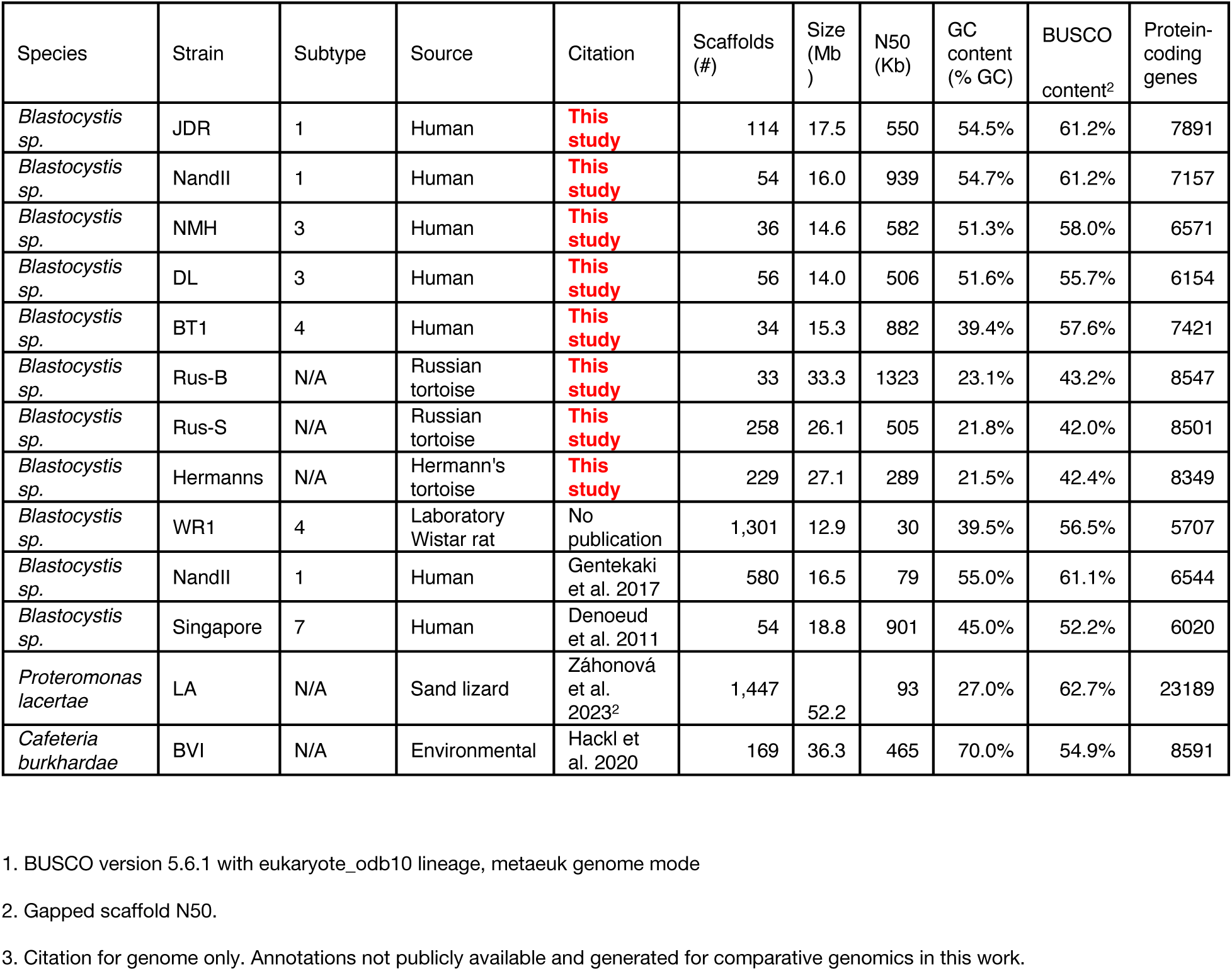
Genome assembly statistics for all genomes used in this study.

Previous genome sequencing studies have raised questions about the ploidy of *Blastocystis* and found evidence of segmentally duplicated blocks of genes. With the highly contiguous genomes and multiple isolates of individual subtypes here, we investigated questions of ploidy and conservation of synteny both within *Blastocystis* subtypes and across the genus. *K*-mer frequency-based analysis using GenomeScope (Ranallo-Benavidez et al. 2020) demonstrated a strong diploid signal in all *Blastocystis* isolates, with limited evidence for aneuploidy or polyploidy. We found that strains within ST3 and within ST1 were highly syntenic (Figure S1). However, synteny between subtypes was much lower, broadly following the relatedness between strains based on 18S sequences.

We used a hybrid method of *de novo* gene prediction and RNA transcript evidence to annotate *Blastocystis* genes (see Methods, Genome assembly and decontamination). Because a substantial fraction of genes in *Blastocystis* ST1 and ST7 do not have DNA-encoded stop codons (Gentekaki et al. 2017; Klimeš et al. 2014) (discussed below), open reading frame based gene predictions from DNA sequence are inaccurate. Leveraging our RNA sequencing data, we were able to generate a comprehensive gene annotation for each *Blastocystis* genome that was able to accurately call genes lacking stop codons (see Methods, Gene annotation). Similar approaches were taken for one previously sequenced ST1 genome (Gentekaki et al. 2017), but published *Blastocystis* genomes for ST4 and ST7 were not annotated with these approaches, leading to incorrect gene annotations. For the *Blastocystis* strains sequenced in this study, the number of genes per *Blastocystis* genome sequenced in this work ranged from 6,154 to 8,547 (Table 1). Protein amino acid identity (AAI) across 753 single-copy orthologs was almost identical between strains of the same subtype (99%-100% AAI for both ST1 and ST3), and decreased to 65-75% between human *Blastocystis* isolates from different subtypes and to 40-42% between human and tortoise isolates (Figure S2). Tortoise isolates had lower pairwise AAI than human isolates, with the most distantly related strains in the tortoise-associated clade having a protein identity of 56% AAI for the same 753 orthologs, suggesting substantial genetic distance within this group.

### Genomic diversity within Blastocystis

When we compared sequenced *Blastocystis* genomes, we found substantial variation in the genomes between human *Blastocystis* isolates and the tortoise-associated clade, including genome size. While the tortoise *Blastocystis* isolate genomes are ∼1.5x-2.4x larger than the human isolate genomes, this difference is not reflected in protein-coding gene content, suggesting that non-genic regions are driving the difference in genome size. We sought to determine why the genomes of *Blastocystis* isolates in the tortoise-associated clade were larger than the genomes of the *Blastocystis* human isolates.

We found little difference in transposable element presence across *Blastocystis*, which ranges from 4.2%-6.7% of the genome in the human isolates and 5.6%-7.5% of the genome in the tortoise isolates. However, we did observe an increase in low-complexity regions in the tortoise isolates, comprising 0.02%-1.4% of the human isolate genomes and 11.8%-12.7% of the tortoise isolate genomes (Table S3). This increase in low complexity regions does not explain the difference in genome size; if these regions are removed, the tortoise isolates remain 1.3x-2.1x larger than the human isolates. These low complexity regions may be related to the overall lower GC content of the tortoise isolates, which ranges from 21-23% in tortoise isolates to 39-55% in human isolates (Table 1).

As repetitive elements did not explain differences in genome size, we investigated differences in other genomic features. We found that the almost all introns in human ST1, ST3, and ST4 *Blastocystis* are exactly 30 bp long, in agreement with previous estimates of intron size in *Blastocystis* ST1 and ST7 (Gentekaki et al. 2017; Denoeud et al. 2011). However, introns in the tortoise isolates are longer and more variable in length (Figure 2A). We additionally found that the intergenic spaces between genes are much longer in the tortoise *Blastocystis* isolate than in human isolates (Figure 2B). The median intergenic length in the tortoise isolates ranges from 645-719 bp, while the median intergenic length in the human isolates ranges from 137-186 bp. Intergenic lengths in the human isolates can frequently be extremely small, only spanning tens of base pairs (Figure 2B). Together, the total amount of DNA spanned by introns and intergenic regions is 13.9-16.5 Mb in the tortoise *Blastocystis* isolates, whereas these regions span only 3.7-5.7 Mb in human *Blastocystis* isolates. Considered together, the larger introns and intergenic distance in the tortoise isolates explain the difference in their genome sizes relative to the human isolate.

**Figure 2.**
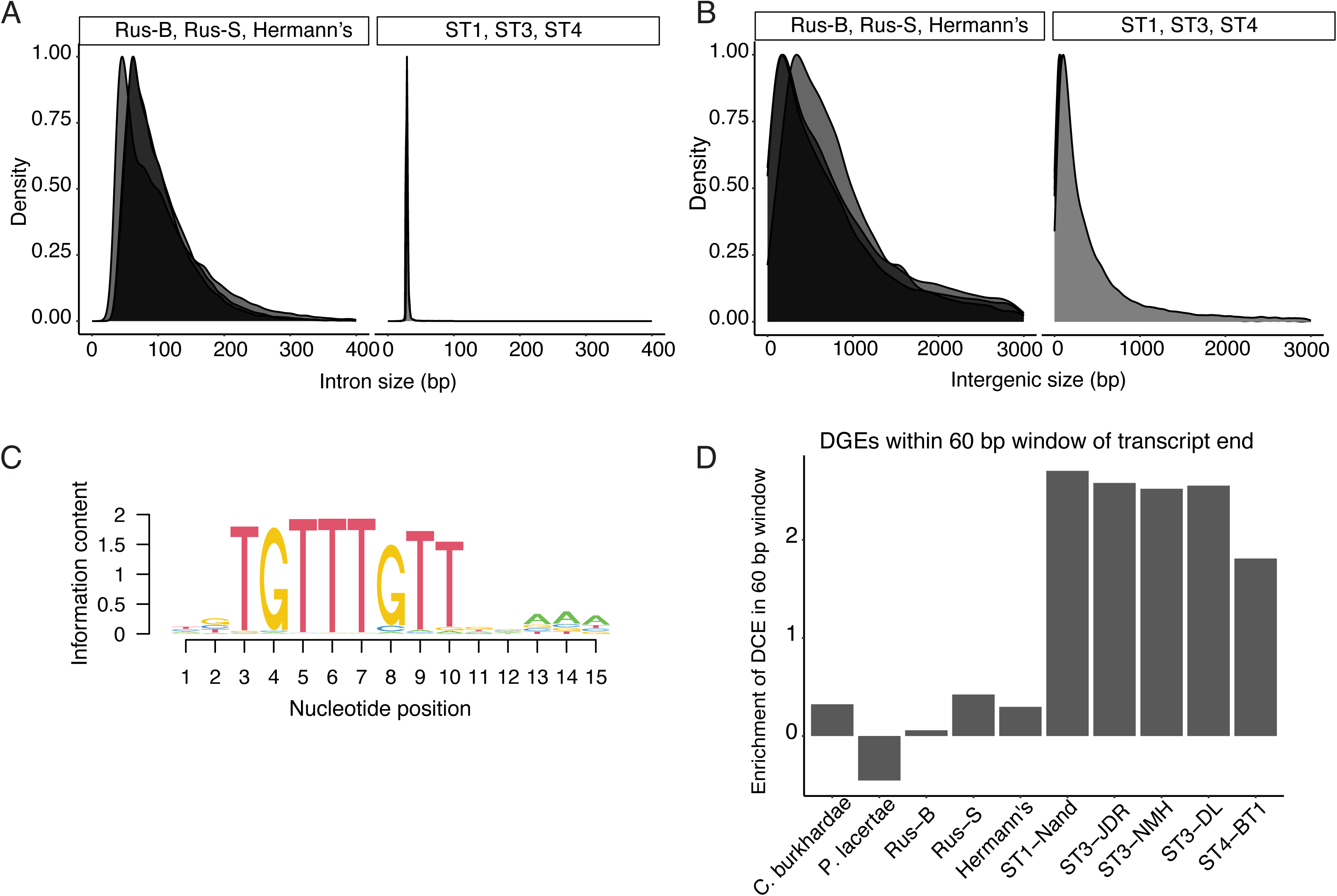
(A) Intron lengths and (B) Intergenic lengths in tortoise isolates (left) and human isolates right). (C) Sequence motif enriched adjacent to transcript ends in ST1, ST3, and ST4. Motif generated with STREME (Bailey 2021). This motif is referred to as “downstream gene element” (DGE). (D) Enrichment (log(observed/expected)) of transcripts with downstream conserved element within 30 bp upstream or downstream of transcript end, normalized by expected DGE occurrence given GC content.

### Loss of stop codons is not universal in Blastocystis

In addition to genome size and gene density, we found surprising differences in how genes are encoded in the genome across *Blastocystis*. In *Blastocystis* ST1 and ST7, a fraction of protein coding genes (15% for ST7 and 26% for ST1) lack canonical DNA-encoded termination codons (Gentekaki et al. 2017; Klimeš et al. 2014). A proposed mechanism for translation of these genes is that the translational machinery instead uses a U or a UG sequence in mRNA, along with one or two adenines from the poly-A tail, to generate UAA and UGA on-the-fly stop codons (Klimeš et al. 2014). Additionally, these genomes have a highly conserved motif (downstream gene element, or DGE) roughly 5 base pairs downstream of transcription stop sites of most genes. These sequences are hypothesized to direct 3’ mRNA cleavage and polyadenylation factors to the pre-mRNA (Gentekaki et al. 2017; Klimeš et al. 2014). In a scenario where the DGE is sufficient for 3’ mRNA cleavage, the loss of DNA-encoded stop codons is driven by relaxation of the selection pressure maintaining them.

We investigated all *Blastocystis* isolates for the DGE motif, finding enrichment downstream of protein coding genes in human-isolated *Blastocystis* strains, but not downstream of tortoise-isolated *Blastocystis* strains or other stramenopiles (Figure 2C-D). We additionally found no evidence of transcripts lacking stop codons in the tortoise-associated *Blastocystis* clade, suggesting that the unique transcriptional cleavage machinery described in *Blastocystis* ST1, ST4, and ST7 (Klimeš et al. 2014; Gentekaki et al. 2017) evolved after the divergence of the tortoise- and the human-associated clades sequenced here.

### Gene losses related to cell morphology and signaling unite the Blastocystis genus

Despite the substantial variation in genome diversity for all *Blastocystis* strains sequenced here, the strains share the typical spherical, non-motile *Blastocystis* cell morphology lacking flagella. We find that all *Blastocystis* isolates are united by substantial gene loss related to this morphological change across multiple functional categories. All *Blastocystis* isolates have lost nearly all cilia- and flagella-related genes (Figure 3, Table S4-5). They have also undergone substantial reduction of dynein and kinesin microtubule motor proteins, retaining what is likely the minimal amount of motor proteins required for intracellular transport. This agrees with recent analyses showing reduction in these gene families in human-associated *Blastocystis* subtypes (Záhonová et al. 2023), and extends that observation by demonstrating that these losses unite the genus. In addition, we find that *Blastocystis* isolates have lost nucleotide cyclases and many genes related to signal transduction (Figure 3, Table S4-5). As cilia are major mediators of cell signaling that use cyclic nucleotides as intermediaries (Johnson and Leroux 2010), the loss of cilia in *Blastocystis* likely corresponds to the loss of nucleotide cyclases and reduction in signal transduction related genes. Thus, our sequencing of diverse *Blastocystis* strains provides strong evidence that the flagella-free morphology and associated gene losses evolved in the common ancestor of the *Blastocystis* genus.

**Figure 3.**
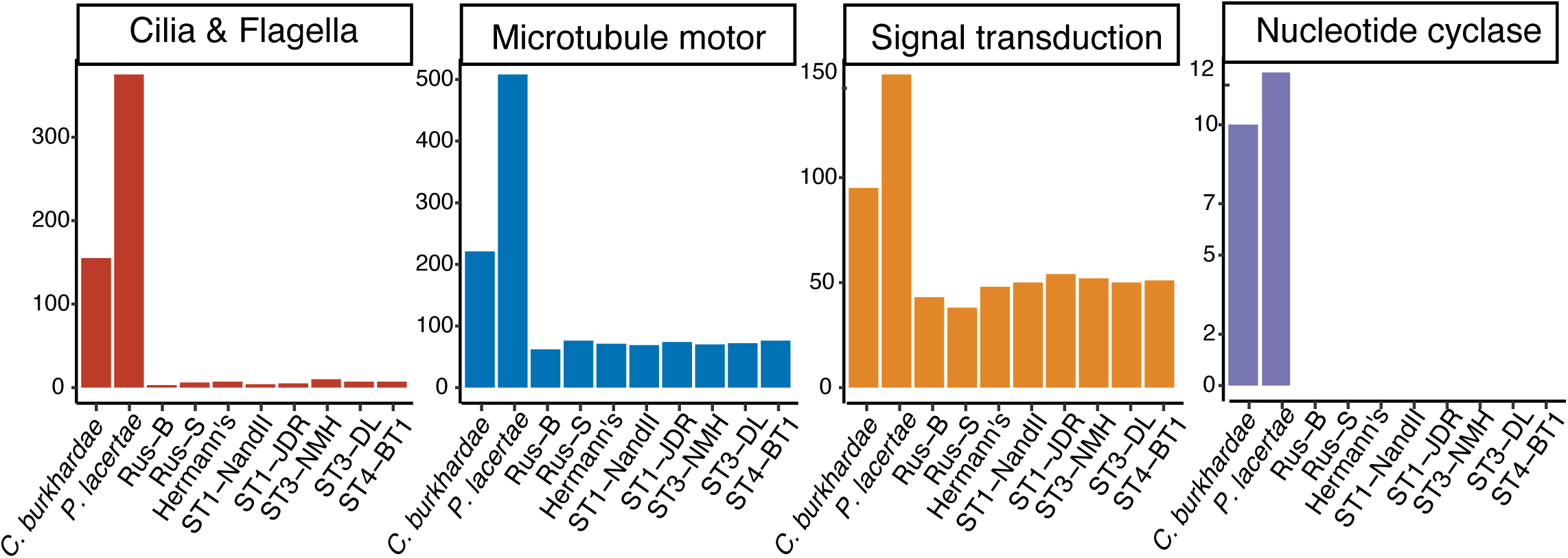
Gene families lost or reduced in all Blastocystis strains. See Table S2 for GO categories and InterPro domains used for defining categories.

### Lineage-specific genes reveal specialization within the Blastocystis genus

We find multiple differences in gene content in between *Blastocystis* isolates that likely reflect their ecological niche. All strains in the herbivorous tortoise-associated *Blastocystis* clade have a larger number of genes involved in plant carbohydrate degradation than the human-associated subtypes. We found five carbohydrate-degrading enzyme families that were either uniquely present in the tortoise isolates or in higher copy numbers, four of which are specific to degrading plant carbohydrates. This includes polygalacturonases, β-D-xylosidases, α-L-arabinofuranosidases, α-L-fucosidases, α-L-rhamnosidases, arabinogalactanases, and β-hexosaminidases (Figure 4). Polygalacturonase is a pectinase that degrades plant cell walls; it was originally characterized in plants, but is also present in phytopathogenic microbes and in bacterial members of ruminant microbiomes (De Lorenzo and Ferrari 2002; Deng et al. 2023). β-D-xylosidases and α-L-arabinofuranosidases are both critical enzymes for degrading xylan, a carbohydrate specific to plant hemicellulose (Yang et al. 2004; Lagaert et al. 2014). The GH154 arabinogalactanase gene family that is present only in the tortoise-associated clade was recently characterized in the gut bacteria *Bacteroides thetaiotamicron* as acting on plant-derived arabinogalactan glycans (Cartmell et al. 2018). Finally, α-L-rhamnosidases are active on a wide variety of polysaccharide residues produced by plants and bacteria (Guillotin et al. 2019). Together, the presence of multiple plant-specific carbohydrate digesting enzymes in the herbivorous tortoise-associated clade strongly suggests that, like the bacterial microbiota, *Blastocystis* utilize dietary plant fiber as a carbon source and participate in the hindgut plant fiber fermentation that is characteristic of herbivorous tortoises (Stevens and Hume 1995).

**Figure 4.**
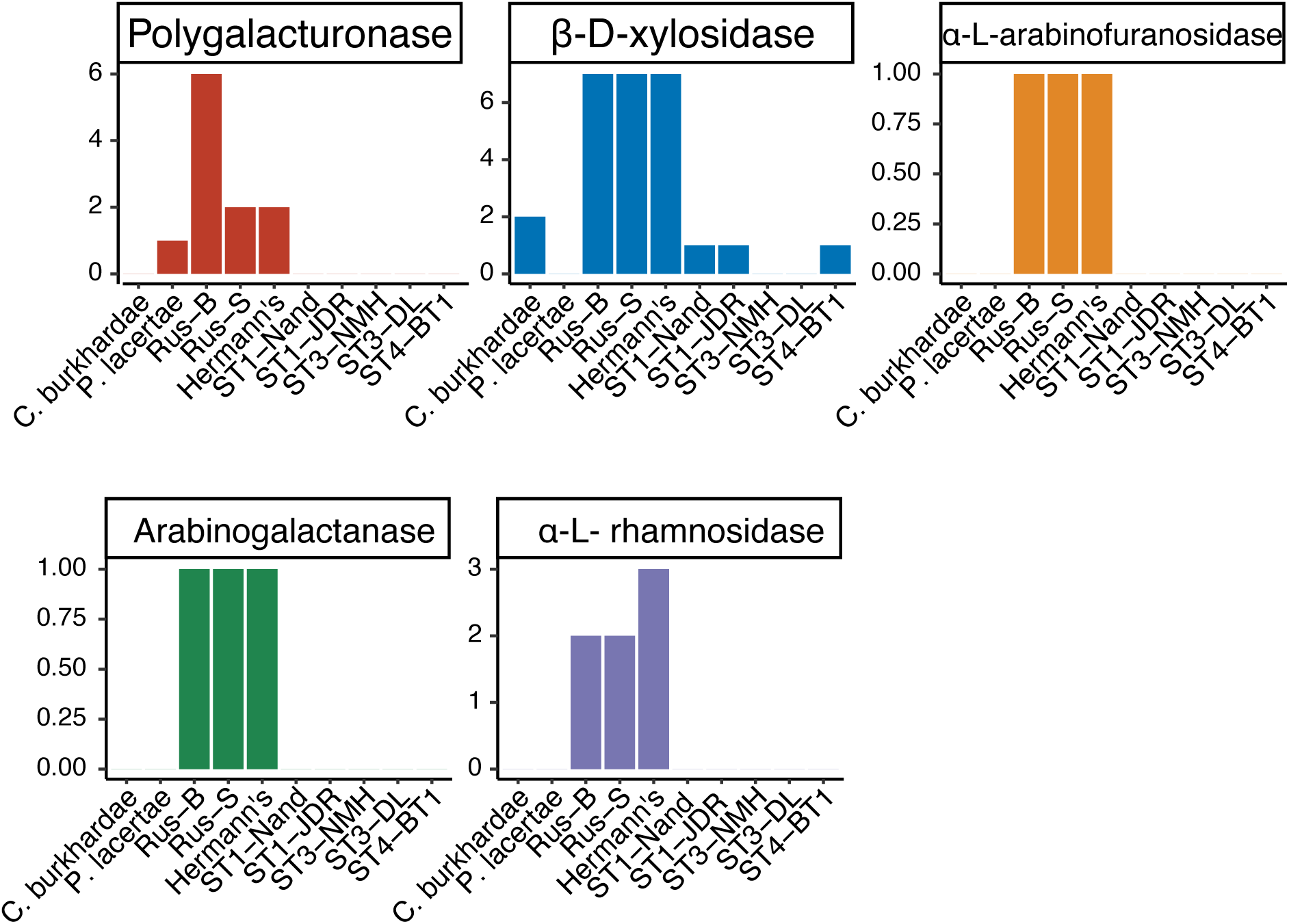
Clade-specific gene family expansions across *Blastocystis,* identified with InterProScan.

### Ongoing horizontal gene transfer from bacteria in the Blastocystis genus

Horizontal gene transfer from bacteria has played an important role in *Blastocystis* evolution, particularly with respect to survival in an anaerobic environment (Eme et al. 2017). With high-quality genomes of *Blastocystis* isolates that span the genus, in combination with the genomes of other stramenopiles, we can estimate times of these transfer events. We re-analyzed HGT events into ST1 previously identified in Eme et al 2017, and additionally screened all sequenced *Blastocystis* genomes for evidence of additional transferred genes (see Methods, Screening for horizontally transferred genes).

Previous analysis of *Blastocystis* ST1, ST4, and ST7 genomes identified 74 horizontal transfer events in *Blastocystis* ST1 from bacteria (Eme et al. 2017). We were able to confidently identify orthologous genes for 67 of these genes (Table S6), of which 27 genes had strong phylogenetic support for horizontal gene transfer from bacteria to eukaryotes, 17 genes lacked strong phylogenetic support for horizontal gene transfer, and the remaining 20 genes were ambiguous. In most ambiguous cases, the *Blastocystis* genes were either sister to an entirely bacterial tree, there were too few orthologs in the NR database to construct a gene tree, or the placing of the *Blastocystis* sequences on the phylogeny lacked strong support.

We additionally screened all genomes sequenced for new cases of HGT. To minimize the impact of potential genome contamination, we limited this screen to genes that were expressed in our RNA-seq data, were shared in more than one *Blastocystis* genome, and were flanked by unambiguously eukaryotic sequence to (see Methods). We found 13 newly identified HGT cases, six which are unique to the tortoise-associated *Blastocystis* lineage.

Interestingly, two of the six tortoise-unique transfers are glycoside hydrolase genes, alpha-L-rhamnosidase and arabinogalactanase, that are uniquely expanded in this lineage and not present in the human *Blastocystis* isolates. The remaining transfers are two GNAT family N-acetyltransferases, a cold-shock protein, and a SMI1/KNR4 family protein (Table S7).

The alpha-L-rhamnosidase gene family cleaves alpha-L-rhamnose from a large diversity of molecules and is present across the tree of life. This gene is present in two copies in Rus-B and Rus-S and three copies in Hermann’s, and is not present in any human *Blastocystis* isolates or in *P. lacertae*. Two genes with the alpha-L-rhamnosidase Interpro domain were present in the *C. burkhardae* genome. We constructed a gene phylogeny of the *C. burkhardae* genes and the tortoise genes together with 588 homologous genes retrieved with a sequence search of the *Blastocystis* and *C. burkhardae* sequences against the clustered NCBI NR database (see Methods). The gene phylogeny was separated into two fully supported clades, one with the *C. burkhardae* sequences, eukaryotic sequences from protists and rotifers, and some bacterial sequences, and the other with bacterial sequences and the *Blastocystis* sequences (Figure 5B). The tortoise sequences grouped together with sequences from gut isolates and gut metagenome-assembled genomes in the Clostridia bacterial class, which contains prevalent members of the animal gut microbiota. Thus, the *Blastocystis* tortoise lineage likely acquired this carbohydrate degrading enzyme from Clostridia bacteria present in the gut microbiome, suggesting a gut-specific function.

**Figure 5.**
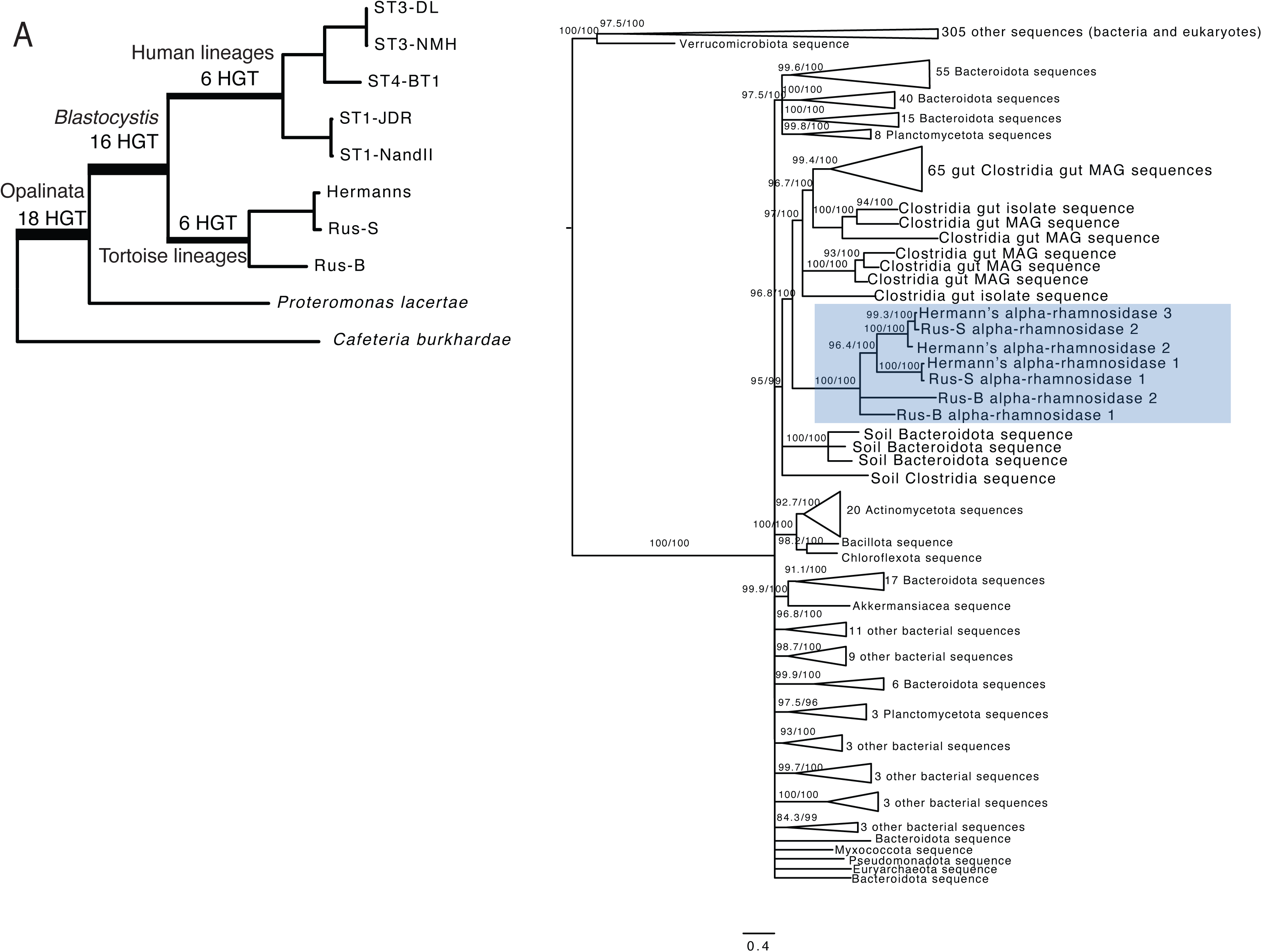
(A) Estimated time of transfer for 40 horizontally transferred genes in *Blastocystis* shared between more than one species or strain. Includes previously identified and validated cases originally identified in (Eme et al. 2017) and newly-identified transfer events. (B) Gene phylogeny of tortoise isolate alpha-rhamnosidase genes and 588 homologous proteins. Support values indicate 1,000 replicates of SH-aLRT test and ultrafast bootstraps (UFboot). Branches with support values <95 for UFboot or <80 for SH-aLRT are collapsed.

The other horizontally transferred carbohydrate gene in the tortoise *Blastocystis* lineage is within the recently described GH154 glycoside hydrolase gene family, characterized in *Bacteroides thetaiotamicron* as acting on plant-derived arabinogalactan glycans (Cartmell et al. 2018). This gene family is present exclusively in the tortoise *Blastocystis* isolates and not found in any other *Blastocystis* isolates or any other stramenopile genomes. A sequence search for the tortoise *Blastocystis* genes against the clustered NCBI NR database did not return any eukaryotic hits. A gene phylogeny of all bacterial homologs and the tortoise *Blastocystis* genes shows the tortoise *Blastocystis* genes group together with bacteria in the *Bacteroidota* (syn. *Bacteroidetes*) phylum (Figure S3).

We did not find strong evidence that the other three carbohydrate gene families uniquely present or expanded in the tortoise *Blastocystis* lineage were horizontally acquired from bacteria. Homologs to the polygalacturonase gene were all eukaryotic. While the evolutionary history of this gene is likely complex as most homologs were within Streptophyta, it was not horizontally acquired from bacteria. β-D-xylosidase and α-L-arabinofuranosidase both appeared to have complex evolutionary histories but without conclusive evidence of horizontal gene transfer. Specifically, several of the β-D-xylosidase genes in the tortoise lineages had no eukaryotic homologs, but a gene phylogeny of all β-D-xylosidase genes including the human isolates did not yield a well-supported branch of the tortoise genes nested within bacteria. Similarly, the α-L-arabinofuranosidase gene had both bacterial and eukaryotic homologs, but there was not strong support for the placement of the *Blastocystis* sequences on this phylogeny either with the eukaryotic sequences or with the bacterial sequences.

Together, these results indicate that horizontal gene transfer played a major role in all evolutionary transitions investigated in this work: at the evolutionary transition from free-living to gut symbiont, in the *Blastocystis* genus broadly, and is an ongoing method of genetic diversification in this organism.

## Discussion

The stramenopile protist *Blastocystis* is the most prevalent protist in the human gut and is widespread throughout vertebrates. Here we present contiguous and complete genomes of eight *Blastocystis* strains, five isolated from humans and three from herbivorous tortoises. Long-read sequencing coupled with metagenomic assembly techniques enabled us to assemble three of these genomes from xenic cultures, which has been technically infeasible with standard short-read approaches. Comparative genomic analyses between these *Blastocystis* genomes alongside other closely related stramenopiles revealed evolutionary changes that the *Blastocystis* genus has undergone to colonize a wide variety of animal guts.

The herbivorous tortoise isolates of *Blastocystis* diverged substantially from the human isolated *Blastocystis,* and have many genomic features not seen in the human isolates. We found substantial changes in genome size, with human isolates possessing significantly shorter introns and intergenic regions than the tortoise isolates, and substantially lower GC content in tortoise isolates. The GC content of the tortoise isolates is substantially lower than the genomes of most sequenced stramenopiles; of the 299 stramenopiles with sequenced genomes deposited in GenBank, only three genomes have lower GC content.

It is not clear whether introns and intergenic regions have shrunk in size after the split of the tortoise-human isolates, or if they have lengthened along the tortoise lineage. Genomic streamlining is a frequently observed process in symbiotic and parasitic organisms (McCutcheon and Moran 2012; Leckenby and Hall 2015) and some parasitic eukaryotes have entirely or almost entirely lost introns (Wang et al. 2018; Wilihoeft et al. 2001). The phenomenon of intergenic and intron size shrinking in *Blastocystis* likely reflects this general process, but the unexpected differences between different host-associated *Blastocystis* isolates highlights that this is an ongoing evolutionary process that can proceed at different rates.

Despite these substantial differences in genomic streamlining, all *Blastocystis* isolates sequenced here have undergone loss of the same genes associated with the more complex flagellated cell morphology of other stramenopiles. *Blastocystis* isolates have lost nearly all cilia- and flagella-related proteins, substantially reduced the number of microtubule motor proteins, and have lost all nucleotide cyclases and many signal transduction related genes. These changes unite all *Blastocystis* isolates, demonstrating shared biology across the genus despite large evolutionary distances and differences in genome structure between isolates.

We additionally find evidence consistent with host adaptation within *Blastocystis*. The *Blastocystis* strain isolated from herbivorous tortoises contained many carbohydrate digesting enzymes not present in other *Blastocystis* isolates (Figure 4). Two of these gene families were acquired via horizontal transfer from bacteria, and in one case via horizontal transfer from bacteria that also colonize the animal gut. The presence of plant-specific carbohydrate digesting enzymes suggests that, like some ciliates and many gut bacteria, *Blastocystis* may ferment plant carbohydrates, a critical element of energy homeostasis in predominantly plant-eating animals (Stevens and Hume 1995).

While *Blastocystis* is widespread across vertebrates, there is limited evidence for co-phylogeny of *Blastocystis* subtypes with their host (Figure 1). Instead, the *Blastocystis* phylogeny more closely resembles that of spore-forming gut bacteria (Moeller et al. 2016), with host switching as a common phenomenon. Because of the lack of strong co-phylogenetic signal, open questions remain about *Blastocystis* host association, including what the barriers to colonizing different hosts are and how multiple strains or subtypes (commonly termed ‘mixed infections’) are maintained, will require a denser sampling of genomes across the *Blastocystis* tree of life. As human associated *Blastocystis* subtypes are not monophyletic, understanding the biology of the protists that impact humans will require considering *Blastocystis* subtypes that are present in the guts of diverse vertebrate hosts, and how these subtypes function in different environments. Finally, the vast differences seen in the phylogenetically distinct tortoise *Blastocystis* isolates sequenced here compared to previous sequenced human-associated *Blastocystis* isolates highlights that substantial genetic diversity in this genus has yet to be explored.

Five of the eight *Blastocystis* strains sequenced in this work were grown in the presence of other microbes. Despite the challenges of generating highly contiguous and contamination free genomes from mixed culture metagenomic sequencing, we successfully combined advances in deep long read DNA sequencing and metagenomic assembly techniques to acquire high-quality genomes from xenic cultures. To address the challenge of annotating genes in genomes where stop codons are not universal, we utilized RNA-sequencing and state-of-the-art annotation pipelines. These two technical accomplishments were essential for enabling robust comparative genomic analyses of Blastocystis and its closest sequenced relatives. The strategy established in this study can be extended in future work to the many eukaryotic microbes that cannot be cultured axenically. The resulting genomes will fill critical gaps in our understanding of eukaryotic microbes and the evolution of this domain of life.

## Methods

### Strain cultivation

*Blastocystis* strains NandII (ATCC 50177), JDR (ATCC 50587), BT1 (ATCC 50608), DL (ATCC 50626), and NMH (ATCC 50754) were obtained from the American Type Culture Collection. The JDR strain was labeled as subtype 3 based on rRNA analysis, but 18S rRNA gene sequencing and subsequent genome sequencing revealed in fact that this strain belongs to subtype 1. All strains were isolated from human sources. Cultures were incubated under anaerobic conditions at 37C on biphasic egg slant media overlaid with Locke’s solution supplemented with 25% heat inactivated horse serum (ATCC Medium 1671). Media was pre-reduced for 24 hours before use. ST1 and ST4 strains were maintained in axenic culture, while ST3 and tortoise isolates were grown in xenic culture alongside other microbes present in the original samples. These xenic culture strains proved resistant to axenization.

*Blastocystis* strains from tortoises were isolated from fresh fecal material that was processed within 8 hours of sampling. Fecal material was resuspended in sterile PBS, strained using a 70µm cell strainer, and centrifuged at 700xg for 5 minutes at room temperature. The pellet was then washed 3 times with PBS before being resuspended in 80% Percoll. This was overlaid with 40% Percoll and centrifuged at 100xg for 10 minutes at room temperature. Protists were removed from the top of the 40% Percoll layer, washed in PBS, and resuspended in pre-reduced growth medium.

### DNA extraction and sequencing

High molecular weight DNA was isolated from *Blastocystis* cultures with a modified CTAB-based protocol (adapted from McLay 2017) that minimizes polysaccharide contamination. *Blastocystis* cells were collected from cultures by pelleting with centrifugation at 800x*g* for 5 minutes and snap frozen at −80° C. Frozen pellets were gently resuspended in two volumes of a high salt CTAB lysis buffer (100 mM Tris pH 8, 1.2 M NaCl, 10 mM EDTA, 4% CTAB, 1% PVP-40). 200 µg/mL RNAse A and 200 µg/mL proteinase K were added before incubation at 60° C for 90 minutes. One volume of phenol:chloroform:isoamyl alcohol (25:24:1) was added and samples were mixed end over end for 15 minutes. Samples were centrifuged at 17000*g* for 10 minutes and the upper aqueous layer was recovered. Two additional extractions were performed with pure chloroform, recovering the upper aqueous layer each time. DNA was next precipitated with 2 volumes of a low salt CTAB precipitation buffer (100 mM Tris pH 8, 10 mM EDTA, 2% CTAB) by end over end rotation at 60° C for 2.5 hours. DNA was pelleted by centrifugation at 17,000x*g* for 20 minutes. DNA pellets were washed twice with 75% ethanol and resuspended in 10 mM Tris pH 8. Genomic DNA was further cleaned with 1.8 volumes of Ampure XP beads according to the manufacturer’s protocol (Beckman Coulter, Brea, CA). RNA was isolated from *Blastocystis* cultures with a Trizol extraction following manufacturer’s protocols (Invitrogen, Waltham, MA).

Nanopore sequencing libraries for all human isolates and for the Rus-B strain were prepared using the Genomic DNA by Ligation protocol SQK-LSK109 (Oxford Nanopore Technologies, Oxford, UK) following the manufacturer’s instructions, sequenced on a full FLO-MIN106D R9 flow cell on a MinION sequencer, and basecalled with guppy version 5.0.16. Nanopore sequencing libraries for the Rus-S and Hermann’s isolates were prepared using the SQK-LSK110 kit LSK109 (Oxford Nanopore Technologies, Oxford, UK) and sequenced on a full R10.4.1 flow cell, and basecalled with dorado version 0.5.3 (https://github.com/nanoporetech/dorado).

Short read Illumina DNA and RNA libraries were prepared using the KAPA HyperPlus PCR-free and KAPA mRNA capture kits (Roche, Basel, Switzerland), respectively, following manufacturer’s instructions. RNA integrity and library fragment size distribution were quantified using an Bioanalyzer 2100 (Agilent, Santa Clara, CA). Illumina DNA and RNA libraries for all human *Blastocystis* isolates and for the Rus-S and Hermann’s tortoise isolates were sequenced on an Illumina NextSeq 500 (Illumina, San Diego, CA), and libraries for the Rus-B tortoise isolate were sequenced on an Illumina MiSeq v3.

### Genome assembly and decontamination

Basecalled nanopore reads were assembled using Flye version 2.9.2 with the -- meta option enabled (Kolmogorov et al. 2019, 2020). Assembly polishing was performed using four rounds of nanopore-read polishing using racon version 1.4.3 (Vaser et al. 2017), three rounds of Illumina DNA polishing using PolyPolish version 0.5.0 (Wick and Holt 2022), and 1 round of short-read polishing using POLCA as implemented in MaSuRCA version 4.0.9 (Zimin et al. 2013).

For the ST4 strain BT1, Hi-C data was generated using Phase Genomics (Seattle, WA) Proximo Hi-C 2.0 Kit, and the Phase Genomics Proximo Hi-C genome scaffolding platform was used to create chromosome scale scaffolds from the hybrid assembled genome. The Hi-C scaffolding resulted only in modest improvements to the original assembly quality; the number of scaffolds reduced from 45 to 34, and the assembly N50 improved to 882 from 876.

We used three different methods to separate eukaryotic, *Blastocystis*-originating sequence from other microbial sequences for xenic and axenic assemblies. For xenic assemblies, sequences were binned using MetaBat2 version 2.17.60 as a binning algorithm, taxonomically annotated using MetaEuk taxtocontig, and additionally annotated as likely eukaryotic or prokaryotic using EukRep (Kang et al. 2019; Levy Karin et al. 2020; West et al. 2018). For binning, short DNA sequencing reads were aligned to the polished assembly using minimap2 version 2.26-r1175 and summarized using the utility jgi_summarize_bam_contig_depths (Li 2018). Metaeuk taxonomy prediction was performed by running version 1f7d7023bdac3d46964ca380bef1b2848685501e, metaeuk easy-predict with mmseqs Swiss-Prot database build and metaeuk taxtocontig with the options ‘--majority 0.5 -- tax-lineage 1 --lca-mode 2’. EukRep v0.6.6. was run in ‘balanced’ mode.

Eukaryotic contigs were identified through a union of MetaEuk contigs annotated as eukaryotic, removing any not identified as eukaryotic by EukRep. We found that MetaEuk was not able to identify *Blastocystis* contigs reliably at lower taxonomic levels than the superkingdom Eukaryota, which we validated by running the pipeline on existing NCBI genomes. Instead, any contigs that were annotated as eukaryotic were considered as potentially *Blastocystis* in origin. We cross-referenced these eukaryotic contigs with metabat bins, finding bins corresponding to eukaryotic sequence. In two cases, eukaryotic sequences that appeared to be *Blastocystis* in origin were split across two metabat bins, and these bins were combined. We removed any sequences from these bins that were annotated by MetaEuk as non-eukaryotic. We verified that these final cleaned bins corresponded to *Blastocystis* by BLASTing the NCBI RefSeq *Blastocystis* coding gene sequences against these bins. In several cases, we found bins that were annotated as eukaryotic but did not have any hits to *Blastocysis* coding sequences. These bins were discarded as likely originating from non-*Blastocystis* eukaryotic organisms in the culture.

We found that contigs containing ribosomal RNA genes were often not binned together with protein coding genes, likely due to their highly repetitive nature and differences in DNA coverage. We searched for contigs containing 18S rRNA genes by BLASTing 18S rRNA genes against the full assembly. Contigs with full length and high percent identity hits to any Blastocystis 18S genes that were also annotated as eukaryotic by MetaEuk were included in final assemblies.

For axenic assemblies, sequenced were not binned with MetaBat2, but MetaEuk and EukRep were performed as described above to identify potential contaminating sequences. This resulted in two contigs being removed from the BT1 assembly, and no contigs being removed from Nand or JDR assemblies.

Genome completeness was determined with sequenced-based and gene-based approaches. For the sequence based approach, we applied a *k*-mer frequency-based approach with GenomeScope v2.0 (Ranallo-Benavidez et al. 2020). Gene-based completeness was determined using BUSCO v5.6.1 with the lineage dataset eukaryote_odb10 in the genome mode (Manni et al. 2021).

### Annotating repetitive elements

Repetitive elements in *Blastocystis* were annotated with a combination of RepeatMasker and RepeatModeler using the Dfam database version 3.8 (Smit et al. 2013; Flynn et al. 2020; Storer et al. 2021). Repetitive elements were predicted from all genomes with RepeatModeler version 2.0.5 with LTRStruct enabled. The predicted transposable elements from all *Blastocystis* genomes that had a putative predicted annotation were combined, and repetitive elements were tabulated in each genome with a combination of the custom repeat library created with RepeatModeler and the Dfam v3.8 database with RepeatMasker 4.1.7-p1.

### Gene annotation

*Blastocystis* genes occasionally do not encode stop codons, which confounds prediction via open reading frame and requires RNA-seq based evidence (Klimeš et al. 2014; Gentekaki et al. 2017). Without consideration of lack of stop codons, common gene annotation errors include insertion of erroneous introns and combining adjacent genes. To accurately annotate the *Blastocystis* genomes, we created a pipeline that used a combination of *de novo* gene prediction with Braker3 and transcriptome assembly, combining all sets of gene models with Mikado (Venturini et al. 2018; Gabriel et al. 2023).

RNA-seq reads were aligned to assemblies with STAR version 2.7.11b (Dobin et al. 2013), with a minimum of 3 supporting reads required for each junction. Transcript assemblies were generated with Stringtie version 2.2.3 (Pertea et al. 2015), which uses RNA-seq alignments against a genome to assemble transcripts. Because *Blastocystis* has very short intergenic regions, transcript assemblers can occasionally combine multiple transcripts into one. To address this, assembled transcripts were split if an exon had zero RNA-seq coverage. Assembled transcripts less than 100 bp long were discarded. To generate *de novo* gene predictions, braker version 3.0.8 was run with OrthoDB11 Eukaryota proteins as protein evidence and the aligned RNA-seq as transcript evidence (Gabriel et al. 2023; Kuznetsov et al. 2022). The Mikado transcript scoring file (available in code repository) was optimized to choose transcripts with strictly RNA-seq supported introns, but to prioritize genes containing stop codons so long as the intron boundaries were strictly supported.

### Functional annotation and orthology inference

Proteins from *Cafeteria burkhardae* BVI (formerly *Cafeteria roenbergensis* BVI), and predicted proteins from *Proteromonas lacertae* were used for comparative genomics analyses (Záhonová et al. 2023; Hackl et al. 2020). The genome assembly, transcripts, and proteins for *C. burkhardae* was downloaded from GenBank under accession GCA_008330645.1. The genome sequence for *Proteromonas lacertae* was downloaded from GenBank under accession GCA_002245135.1. As no predicted proteins or RNA-seq data was available for *P. lacertae*, genes were predicted from this genome using the Braker3 pipeline, using OrthoDB11 Eukaryota protein sequences as evidence. Proteins from all *Blastocystis* assemblies, *C. burkhardae*, and *P. lacertae* were annotated using a combination of InterProScan version 5.66 (Jones et al. 2014), DeepLoc2 to predict protein localization (Thumuluri et al. 2022), and dbCAN version 4.0.0 to annotate carbohydrate active enzymes (Zhang et al. 2018). Orthologous sequences were determined using OrthoFinder2 with diamond as the search algorithm and mcl as the clustering algorithm (Emms and Kelly 2019; Buchfink et al. 2015; Li et al. 2003).

Genes related to cilia and flagella, microtubule motor proteins, signal transduction, ion channels, and nucleotide cyclase were identified using either GO categories or Interpro domains corresponding to these categories, and are listed in Table S4. Carbohydrate-degrading enzymes were identified based on the gene annotation methods mentioned above; specifically, polygalacturonase enzymes were identified with the PANTHER domain PTHR33928, beta-D-xylosidase with the Interpro domain IPR044993, a-L-arabinofuranosidase with the Interpro domain IPR010720, a-L-rhamnosidase with the PANTHER domain PTHR34987, and the GH154 arabinogalactanase with the PANTHER domain PTHR35339.

### Identifying motifs adjacent to transcript ends

To identify motifs enriched downstream of transcript ends, we extracted a 60 bp window up and downstream of all annotated transcript ends in all *Blastocystis* genomes sequenced here, as well as *P. lacertae* and *C. burkhardae*. Motifs were identified separately in each genome using STREME version 5.5.7 (Bailey 2021). A motif similar to the one identified by Klimeš et al 2014 was identified in both ST1, both ST3, and the ST4 human isolate, but no significant and informative motifs were identified in the tortoise-isolated *Blastocystis* isolates or in the *P. lacertae* or *C. burkhardae* genomes. We then combined the sequences from all human isolates and ran STREME on this combined dataset to generate a sequence motif (Figure 2).

To determine enrichment of this motif in all genomes, we searched all transcript ends for matches to a degenerate version of the core region of the sequence motif, TGTTGTT, allowing up to two mismatches. As this motif is inherently an AT-rich sequence, the probability of observing it by chance is a function of the GC content of the sequence that is searched. We calculated the expected probability of seeing the 7-base pair in each genome given the transcript window GC content and the number of transcripts, and calculated the enrichment of this motif as the log of the observed motif counts divided by the background expectation value (Figure 2).

### Strain and species phylogenies

The *Blastocystis* genus phylogeny was constructed using 18S rRNA sequences deposited in GenBank (Table S1) along with 18S rRNA sequences extracted from each *Blastocystis* assembly and from the *Proteromonas lacertae* genome as an outgroup. Sequences were aligned with MAFFT v7.505 (Katoh and Standley 2013) and a maximum likelihood phylogeny constructed using IQ-TREE version 2.3.5 (Minh et al. 2020) using with the GTR sequence model, 1,000 ultrafast bootstrap and SH-aLRT replicates, and rooted on *P. lacertae*.

To construct a species phylogeny of all species and strains with genomic data used in this study, 13 conserved protein-coding genes identified with BUSCO version 5.6.1 as described above were extracted from *Cafeteria burkhardae, Proteromonas lacertae*, and all *Blastocystis* strains sequenced here. Proteins were aligned individually with MAFFT v7.505 and concatenated. Best-fit models for each partition were selected using a relaxed hierarchical clustering algorithm as implemented by ModelFinder in IQ-TREE version 2.3.5 (Chernomor et al. 2016; Minh et al. 2020; Kalyaanamoorthy et al. 2017). 1,000 ultrafast bootstrap and SH-aLRT replicates were performed.

### Gene contamination and horizontal gene transfer screens

*Blastocystis* genomes were screened for bacterial contamination and for possible horizontal gene transfer events using a modified version of an alien index score (Gladyshev et al. 2008; Wisecaver et al. 2016). The alien index score compares the best bit score of a query sequence to a group of related taxa versus to all other unrelated groups, and normalizes this score by the maximum possible bit score of the query sequence. It is determined by the formula 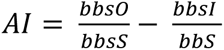, where bbsO is the bit score of the best hit to a species outside of the group lineage, bbsI is the bit score of the best hit to a species within the group lineage (skipping self hits), and bbsS is the bit score of the query aligned to itself. The alien index was calculated for each *Blastocystis* protein by querying proteins against the NCBI nr database using DIAMOND version 2.1.6 (Buchfink et al. 2015)(code availability: https://github.com/allind/alienindex). The self-group was designated as *Blastocystis* (taxid: 12967), and the group lineage was set as all eukaryotes (taxid: 2759).

As an additional control for whether our genomes had high levels of genome contamination, we determined what fraction of genes with high alien index scores were likely transcribed by *Blastocystis.* Our RNA-seq strategy involved poly-A purification, which removes non-polyadenylated transcripts. As polyadenylation is unique to eukaryotes, this removes bacterial transcripts. We quantified gene expression using kallisto version v0.48.0 (Bray et al. 2016), and determined whether genes with alien index scores greater than 0.2 were expressed and therefore likely truly integrated into the *Blastocystis* genomes. The number of genes with alien index scores above 0.2 ranged from 59-81 genes across all sequenced genomes, and greater than 85% were expressed in all genomes. Genes with alien index scores greater than 0.2 that were expressed at TPM > 1 and had orthologs in one or more other *Blastocystis* genomes were further examined for evidence of horizontal gene transfer with gene tree reconstruction.

### Gene tree reconstruction for horizontal transfer

Phylogenetic trees for putative horizontal gene transfer candidates were constructed from all diamond blastp hits using the NCBI NR database clustered at 90% sequence identity and 90% sequence length (Buchfink et al. 2015). Sequence hits that were outliers in length were excluded from downstream analyses. Protein sequences were aligned with MAFFT v7.505 (Katoh and Standley 2013), and maximum likelihood phylogenies were constructed using IQ-TREE version 2.3.5 using ModelFinder to determine the best-fit substitution model, and with 1,000 ultrafast bootstrap replicates and 1,000 replicates of the SH-like approximate likelihood ratio test (Guindon et al. 2010). Branches with SH-aLRT support less than 70 and UF-boot less than 80 were collapsed and trees were midpoint rooted before being examined for horizontal transfer.

## Acknowledgements

We thank members of the Pollard lab and the Bhatt lab for helpful discussions. This work was supported NHLBI grant#R01-HL160862, Chan Zuckberg Biohub SF and Gladstone Institutes to K.S.P, by the Benioff Center for Microbiome Medicine Trainee Pilot Award and Georgia Institute of Technology to A.L.L. A.SB. is supported by NIH R01AI143757 and R01AI148623, as well as a Stand Up 2 Cancer grant and a Paul Allen Distinguished Investigator Award.

## Author Contributions

Conceptualization – A.L.L, A.S.B, K.S.P.; Methodology – A.L.L., N.A.M, E.S.G, A.S.B, K.S.P.; Investigation – A.L.L, K.S.P.; Resources – A.S.B., K.S.P.; Writing – original draft – A.L.L., N.A.M, E.S.G, A.S.B, K.S.P; Writing – review & editing - A.L.L., N.A.M, E.S.G, A.S.B, K.S.P; Visualization – A.L.L.; Supervision – K.S.P., A.S.B.; Funding acquisition – A.L.L., K.S.P., A.S.B.

## DATA ACCESS

Genome assemblies and annotations are available on Genbank (Table 2) and are additionally available on Figshare at https://figshare.com/projects/Blastocystis_genomes/186136. As no gene annotations were available for *Proteromonas lacertae*, BRAKER3 predictions were generated and have been made publicly available in this repository as well. RNA sequencing for all strains is available from the SRA under BioProject PRJNA1041441. Code to calculate the alien index is available at https://github.com/allind/alienindex. All other code used in this study is available at https://github.com/allind/blastannotate. All alignments and phylogenies generated in this work are available at https://doi.org/10.6084/m9.figshare.28143281.v1.

**Table 2.** NCBI accessions for genomes generated here.

| Strain | NCBI Bioproject |
| --- | --- |
| Rus-B (also Blastoise) | PRJNA1084774 |
| Rus-S | Processing on Genbank, currently available on Figshare |
| Hermann's | Processing on Genbank, currently available on Figshare |
| ST1-NandII | PRJNA1084796 |
| ST1-JDR | PRJNA1084798 |
| ST3-DL | PRJNA1084795 |
| ST3-NMH | PRJNA1084799 |
| ST4-BT1 | PRJNA1084775 |

---

Figure S1. Synteny of *Blastocystis* genomes based on reciprocal best blast hits. Plot created with ODP (Schultz et al. 2023).

Figure S2. Median percent identity across 753 single-copy orthologs in *Blastocystis*.

Figure S3. Gene phylogeny of tortoise Blastocystis GH154 arabinogalactanase enzyme. Support values indicate 1,000 replicates of SH-aLRT test and ultrafast bootstrap. Branches with aLRT support <80 and UFboot support <90 are collapsed. Tree is midpoint rooted.

Table S1. GenBank accessions of 18S rRNA sequences used to construct *Blastocystis* phylogeny.

Table S2. Amount and percentages of genome masked per strain by RepeatMasker.

Table S3. GO and IPR categories used for gene loss described in Figure 3.

Table S4. Genes containing InterPro domains per genome assembly.

Table S5. Percentage of high alien index genes per genome with poly-A purified RNA-seq expression support.

Table S6. Validation of previously reported ST1-NandII horizontal gene transfers.

Table S7. Newly identified HGT events in *Blastocystis*.

## References

Asnicar F, Berry SE, Valdes AM, Nguyen LH, Piccinno G, Drew DA, Leeming E, Gibson R, Le Roy C, Khatib HA, et al. 2021. Microbiome connections with host metabolism and habitual diet from 1,098 deeply phenotyped individuals. Nat Med 27: 321–332.

Bailey TL. 2021. STREME: accurate and versatile sequence motif discovery. Bioinforma Oxf Engl 37: 2834–2840.

Beghini F, Pasolli E, Truong TD, Putignani L, Cacciò SM, Segata N. 2017. Large-scale comparative metagenomics of Blastocystis, a common member of the human gut microbiome. ISME J 11: 2848–2863.

Boreham PFL, Stenzel DJ. 1993. Blastocystis in Humans and Animals: Morphology, Biology, and Epizootiology. In Advances in Parasitology (eds. J.R. Baker and R. Muller), Vol. 32 of, pp. 1–70, Academic Press https://www.sciencedirect.com/science/article/pii/S0065308X08602067 (Accessed October 24, 2023).

Bray NL, Pimentel H, Melsted P, Pachter L. 2016. Near-optimal probabilistic RNA-seq quantification. Nat Biotechnol 34: 525–527.

Buchfink B, Xie C, Huson DH. 2015. Fast and sensitive protein alignment using DIAMOND. Nat Methods 12: 59–60.

Campo J del, Bass D, Keeling PJ. 2020. The eukaryome: Diversity and role of microeukaryotic organisms associated with animal hosts. Funct Ecol 34: 2045–2054.

Cartmell A, Muñoz-Muñoz J, Briggs JA, Ndeh DA, Lowe EC, Baslé A, Terrapon N, Stott K, Heunis T, Gray J, et al. 2018. A surface endogalactanase in Bacteroides thetaiotaomicron confers keystone status for arabinogalactan degradation. Nat Microbiol 3: 1314–1326.

Chernomor O, von Haeseler A, Minh BQ. 2016. Terrace Aware Data Structure for Phylogenomic Inference from Supermatrices. Syst Biol 65: 997–1008.

David LA, Maurice CF, Carmody RN, Gootenberg DB, Button JE, Wolfe BE, Ling AV, Devlin AS, Varma Y, Fischbach MA, et al. 2014. Diet rapidly and reproducibly alters the human gut microbiome. Nature 505: 559–563.

De Lorenzo G, Ferrari S. 2002. Polygalacturonase-inhibiting proteins in defense against phytopathogenic fungi. Curr Opin Plant Biol 5: 295–299.

Deng Q, Sun X, Gao D, Wang Y, Liu Y, Li N, Wang Z, Liu M, Wang J, Wang Q. 2023. Characterization of Two Novel Rumen-Derived Exo-Polygalacturonases: Catalysis and Molecular Simulations. Microorganisms 11: 760.

Denoeud F, Roussel M, Noel B, Wawrzyniak I, Da Silva C, Diogon M, Viscogliosi E, Brochier-Armanet C, Couloux A, Poulain J, et al. 2011. Genome sequence of the stramenopile Blastocystis, a human anaerobic parasite. Genome Biol 12: R29.

Dobin A, Davis CA, Schlesinger F, Drenkow J, Zaleski C, Jha S, Batut P, Chaisson M, Gingeras TR. 2013. STAR: ultrafast universal RNA-seq aligner. Bioinforma Oxf Engl 29: 15–21.

Eme L, Gentekaki E, Curtis B, Archibald JM, Roger AJ. 2017. Lateral Gene Transfer in the Adaptation of the Anaerobic Parasite Blastocystis to the Gut. Curr Biol CB 27: 807–820.

Emms DM, Kelly S. 2019. OrthoFinder: phylogenetic orthology inference for comparative genomics. Genome Biol 20: 238.

Flynn JM, Hubley R, Goubert C, Rosen J, Clark AG, Feschotte C, Smit AF. 2020. RepeatModeler2 for automated genomic discovery of transposable element families. Proc Natl Acad Sci 117: 9451–9457.

Gabriel L, Brůna T, Hoff KJ, Ebel M, Lomsadze A, Borodovsky M, Stanke M. 2023. BRAKER3: Fully Automated Genome Annotation Using RNA-Seq and Protein Evidence with GeneMark-ETP, AUGUSTUS and TSEBRA. BioRxiv Prepr Serv Biol 2023.06.10.544449.

Gantois N, Lamot A, Seesao Y, Creusy C, Li L-L, Monchy S, Benamrouz-Vanneste S, Karpouzopoulos J, Bourgain J-L, Rault C, et al. 2020. First Report on the Prevalence and Subtype Distribution of Blastocystis sp. in Edible Marine Fish and Marine Mammals: A Large Scale-Study Conducted in Atlantic Northeast and on the Coasts of Northern France. Microorganisms 8: 460.

Gentekaki E, Curtis BA, Stairs CW, Klimeš V, Eliáš M, Salas-Leiva DE, Herman EK, Eme L, Arias MC, Henrissat B, et al. 2017. Extreme genome diversity in the hyper-prevalent parasitic eukaryote Blastocystis. PLOS Biol 15: e2003769.

Godoy-Vitorino F, Goldfarb KC, Karaoz U, Leal S, Garcia-Amado MA, Hugenholtz P, Tringe SG, Brodie EL, Dominguez-Bello MG. 2012. Comparative analyses of foregut and hindgut bacterial communities in hoatzins and cows. ISME J 6: 531–541.

Gough R, Barratt J, Stark D, Ellis J. 2020. Diversity profiling of xenic cultures of Dientamoeba fragilis following systematic antibiotic treatment and prospects for genome sequencing. Parasitology 147: 29–38.

Guillotin L, Kim H, Traore Y, Moreau P, Lafite P, Coquoin V, Nuccio S, de Vaumas R, Daniellou R. 2019. Biochemical Characterization of the α-l-Rhamnosidase DtRha from Dictyoglomus thermophilum: Application to the Selective Derhamnosylation of Natural Flavonoids. ACS Omega 4: 1916–1922.

Guindon S, Dufayard J-F, Lefort V, Anisimova M, Hordijk W, Gascuel O. 2010. New Algorithms and Methods to Estimate Maximum-Likelihood Phylogenies: Assessing the Performance of PhyML 3.0. Syst Biol 59: 307–321.

Hackl T, Martin R, Barenhoff K, Duponchel S, Heider D, Fischer MG. 2020. Four high-quality draft genome assemblies of the marine heterotrophic nanoflagellate Cafeteria roenbergensis. Sci Data 7: 29.

Hoang DT, Chernomor O, von Haeseler A, Minh BQ, Vinh LS. 2018. UFBoot2: Improving the Ultrafast Bootstrap Approximation. Mol Biol Evol 35: 518–522.

Jinatham V, Maxamhud S, Popluechai S, Tsaousis AD, Gentekaki E. 2021. Blastocystis One Health Approach in a Rural Community of Northern Thailand: Prevalence, Subtypes and Novel Transmission Routes. Front Microbiol 12. https://www.frontiersin.org/journals/microbiology/articles/10.3389/fmicb.2021.746340/full (Accessed December 13, 2024).

Johnson J-LF, Leroux MR. 2010. cAMP and cGMP signaling: sensory systems with prokaryotic roots adopted by eukaryotic cilia. Trends Cell Biol 20: 435–444.

Jones P, Binns D, Chang H-Y, Fraser M, Li W, McAnulla C, McWilliam H, Maslen J, Mitchell A, Nuka G, et al. 2014. InterProScan 5: genome-scale protein function classification. Bioinformatics 30: 1236–1240.

Kalyaanamoorthy S, Minh BQ, Wong TKF, von Haeseler A, Jermiin LS. 2017. ModelFinder: fast model selection for accurate phylogenetic estimates. Nat Methods 14: 587–589.

Kang DD, Li F, Kirton E, Thomas A, Egan R, An H, Wang Z. 2019. MetaBAT 2: an adaptive binning algorithm for robust and efficient genome reconstruction from metagenome assemblies. PeerJ 7: e7359.

Katoh K, Standley DM. 2013. MAFFT Multiple Sequence Alignment Software Version 7: Improvements in Performance and Usability. Mol Biol Evol 30: 772–780.

Keeling PJ, Burki F, Wilcox HM, Allam B, Allen EE, Amaral-Zettler LA, Armbrust EV, Archibald JM, Bharti AK, Bell CJ, et al. 2014. The Marine Microbial Eukaryote Transcriptome Sequencing Project (MMETSP): Illuminating the Functional Diversity of Eukaryotic Life in the Oceans through Transcriptome Sequencing. PLOS Biol 12: e1001889.

Khaled S, Gantois N, Ly AT, Senghor S, Even G, Dautel E, Dejager R, Sawant M, Baydoun M, Benamrouz-Vanneste S, et al. 2020. Prevalence and Subtype Distribution of Blastocystis sp. in Senegalese School Children. Microorganisms 8: 1408.

Klimeš V, Gentekaki E, Roger AJ, Eliáš M. 2014. A Large Number of Nuclear Genes in the Human Parasite Blastocystis Require mRNA Polyadenylation to Create Functional Termination Codons. Genome Biol Evol 6: 1956–1961.

Kolmogorov M, Bickhart DM, Behsaz B, Gurevich A, Rayko M, Shin SB, Kuhn K, Yuan J, Polevikov E, Smith TPL, et al. 2020. metaFlye: scalable long-read metagenome assembly using repeat graphs. Nat Methods 17: 1103–1110.

Kolmogorov M, Yuan J, Lin Y, Pevzner PA. 2019. Assembly of long, error-prone reads using repeat graphs. Nat Biotechnol 37: 540–546.

Kostka M. 2017. Opalinata. In Handbook of the Protists (eds. J.M. Archibald, A.G.B. Simpson, and C.H. Slamovits), pp. 543–565, Springer International Publishing, Cham 10.1007/978-3-319-28149-0_4 (Accessed July 28, 2023).

Kuznetsov D, Tegenfeldt F, Manni M, Seppey M, Berkeley M, Kriventseva EV, Zdobnov EM. 2022. OrthoDB v11: annotation of orthologs in the widest sampling of organismal diversity. Nucleic Acids Res 51: D445–D451.

Lagaert S, Pollet A, Courtin CM, Volckaert G. 2014. β-Xylosidases and α-l-arabinofuranosidases: Accessory enzymes for arabinoxylan degradation. Biotechnol Adv 32: 316–332.

Leckenby A, Hall N. 2015. Genomic changes during evolution of animal parasitism in eukaryotes. Curr Opin Genet Dev 35: 86–92.

Levy Karin E, Mirdita M, Söding J. 2020. MetaEuk—sensitive, high-throughput gene discovery, and annotation for large-scale eukaryotic metagenomics. Microbiome 8: 48.

Li H. 2018. Minimap2: pairwise alignment for nucleotide sequences. Bioinformatics 34: 3094– 3100.

Li L, Stoeckert CJ, Roos DS. 2003. OrthoMCL: identification of ortholog groups for eukaryotic genomes. Genome Res 13: 2178–89.

Manni M, Berkeley MR, Seppey M, Simão FA, Zdobnov EM. 2021. BUSCO Update: Novel and Streamlined Workflows along with Broader and Deeper Phylogenetic Coverage for Scoring of Eukaryotic, Prokaryotic, and Viral Genomes. Mol Biol Evol 38: 4647–4654.

McCutcheon JP, Moran NA. 2012. Extreme genome reduction in symbiotic bacteria. Nat Rev Microbiol 10: 13–26.

McLay T. 2017. High quality DNA extraction protocol from recalcitrant plant tissues v1. https://www.protocols.io/view/high-quality-dna-extraction-protocol-from-recalcit-i8jchun (Accessed January 3, 2025).

Minh BQ, Schmidt HA, Chernomor O, Schrempf D, Woodhams MD, von Haeseler A, Lanfear R. 2020. IQ-TREE 2: New Models and Efficient Methods for Phylogenetic Inference in the Genomic Era. Mol Biol Evol 37: 1530–1534.

Moeller AH, Caro-Quintero A, Mjungu D, Georgiev AV, Lonsdorf EV, Muller MN, Pusey AE, Peeters M, Hahn BH, Ochman H. 2016. Cospeciation of gut microbiota with hominids. Science 353: 380–382.

Moss EL, Maghini DG, Bhatt AS. 2020. Complete, closed bacterial genomes from microbiomes using nanopore sequencing. Nat Biotechnol 38: 701–707.

Nash AK, Auchtung TA, Wong MC, Smith DP, Gesell JR, Ross MC, Stewart CJ, Metcalf GA, Muzny DM, Gibbs RA, et al. 2017. The gut mycobiome of the Human Microbiome Project healthy cohort. Microbiome 5: 153.

Nieves-Ramírez ME, Partida-Rodríguez O, Laforest-Lapointe I, Reynolds LA, Brown EM, Valdez-Salazar A, Morán-Silva P, Rojas-Velázquez L, Morien E, Parfrey LW, et al. 2018. Asymptomatic Intestinal Colonization with Protist Blastocystis Is Strongly Associated with Distinct Microbiome Ecological Patterns. mSystems 3.

Pertea M, Pertea GM, Antonescu CM, Chang T-C, Mendell JT, Salzberg SL. 2015. StringTie enables improved reconstruction of a transcriptome from RNA-seq reads. Nat Biotechnol 33: 290–295.

Piperni E, Nguyen LH, Manghi P, Kim H, Pasolli E, Andreu-Sánchez S, Arrè A, Bermingham KM, Blanco-Míguez A, Manara S, et al. 2024. Intestinal Blastocystis is linked to healthier diets and more favorable cardiometabolic outcomes in 56,989 individuals from 32 countries. Cell 187: 4554–4570.e18.

Ranallo-Benavidez TR, Jaron KS, Schatz MC. 2020. GenomeScope 2.0 and Smudgeplot for reference-free profiling of polyploid genomes. Nat Commun 11: 1432.

Saary P, Mitchell AL, Finn RD. 2020. Estimating the quality of eukaryotic genomes recovered from metagenomic analysis with EukCC. Genome Biol 21: 244.

Santin M, Figueiredo A, Molokin A, George NS, Köster PC, Dashti A, González-Barrio D, Carmena D, Maloney JG. 2024. Division of Blastocystis ST10 into three new subtypes: ST42-ST44. J Eukaryot Microbiol 71: e12998.

Scanlan PD, Marchesi JR. 2008. Micro-eukaryotic diversity of the human distal gut microbiota: qualitative assessment using culture-dependent and -independent analysis of faeces. ISME J 2: 1183–1193.

Scanlan PD, Stensvold CR. 2013. Blastocystis: getting to grips with our guileful guest. Trends Parasitol 29: 523–529.

Scanlan PD, Stensvold CR, Rajilić-Stojanović M, Heilig HGHJ, De Vos WM, O’Toole PW, Cotter PD. 2014. The microbial eukaryote Blastocystis is a prevalent and diverse member of the healthy human gut microbiota. FEMS Microbiol Ecol 90: 326–330.

Schultz DT, Haddock SHD, Bredeson JV, Green RE, Simakov O, Rokhsar DS. 2023. Ancient gene linkages support ctenophores as sister to other animals. Nature 618: 110–117.

Šimková H, Câmara AS, Mascher M. 2024. Hi-C techniques: from genome assemblies to transcription regulation. J Exp Bot 75: 5357–5365.

Smit A, Hubley R, Green P. 2013. Repeatmasker Open-4.0. http://www.repeatmasker.org (Accessed January 10, 2015).

Sommer F, Bäckhed F. 2013. The gut microbiota--masters of host development and physiology. Nat Rev Microbiol 11: 227–238.

Stensvold CR, Clark CG. 2020. Pre-empting Pandora’s Box: Blastocystis Subtypes Revisited. Trends Parasitol 36: 229–232.

Stensvold CR, Suresh GK, Tan KSW, Thompson RCA, Traub RJ, Viscogliosi E, Yoshikawa H, Clark CG. 2007. Terminology for Blastocystis subtypes--a consensus. Trends Parasitol 23: 93–96.

Stevens CE, Hume ID. 1995. Comparative physiology of the vertebrate digestive system. 2nd ed. Cambridge University Press, Cambridge http://catdir.loc.gov/catdir/toc/cam026/95013969.html (Accessed October 16, 2023).

Storer J, Hubley R, Rosen J, Wheeler TJ, Smit AF. 2021. The Dfam community resource of transposable element families, sequence models, and genome annotations. Mob DNA 12: 2.

Tan KSW. 2008. New Insights on Classification, Identification, and Clinical Relevance of Blastocystis spp. Clin Microbiol Rev 21: 639–665.

Thorman AW, Adkins G, Conrey SC, Burrell AR, Yu Y, White B, Burke R, Haslam D, Payne DC, Staat MA, et al. 2023. Gut Microbiome Composition and Metabolic Capacity Differ by FUT2 Secretor Status in Exclusively Breastfed Infants. Nutrients 15: 471.

Thumuluri V, Almagro Armenteros JJ, Johansen AR, Nielsen H, Winther O. 2022. DeepLoc 2.0: multi-label subcellular localization prediction using protein language models. Nucleic Acids Res 50: W228–W234.

Vaser R, Sović I, Nagarajan N, Šikić M. 2017. Fast and accurate de novo genome assembly from long uncorrected reads. Genome Res 27: 737–746.

Wang SE, Amir AS, Nguyen T, Poole AM, Simoes-Barbosa A. 2018. Spliceosomal introns in Trichomonas vaginalis revisited. Parasit Vectors 11: 607.

West PT, Probst AJ, Grigoriev IV, Thomas BC, Banfield JF. 2018. Genome-reconstruction for eukaryotes from complex natural microbial communities. Genome Res 28: 569–580.

Wick RR, Holt KE. 2022. Polypolish: Short-read polishing of long-read bacterial genome assemblies. PLOS Comput Biol 18: e1009802.

Wilihoeft U, Campos-Góngora E, Touzni S, Bruchhaus I, Tannich E. 2001. Introns of Entamoeba histolytica and Entamoeba dispar. Protist 152: 149–156.

Yadav M, Verma MK, Chauhan NS. 2018. A review of metabolic potential of human gut microbiome in human nutrition. Arch Microbiol 200: 203–217.

Yang JK, Yoon HJ, Ahn HJ, Lee BI, Pedelacq J-D, Liong EC, Berendzen J, Laivenieks M, Vieille C, Zeikus GJ, et al. 2004. Crystal structure of beta-D-xylosidase from Thermoanaerobacterium saccharolyticum, a family 39 glycoside hydrolase. J Mol Biol 335: 155–165.

Yoshikawa H, Koyama Y, Tsuchiya E, Takami K. 2016. Blastocystis phylogeny among various isolates from humans to insects. Parasitol Int 65: 750–759.

Záhonová K, Low RS, Warren CJ, Cantoni D, Herman EK, Yiangou L, Ribeiro CA, Phanprasert Y, Brown IR, Rueckert S, et al. 2023. Evolutionary analysis of cellular reduction and anaerobicity in the hyper-prevalent gut microbe Blastocystis. Curr Biol CB 33: 2449–2464.e8.

Zhang H, Yohe T, Huang L, Entwistle S, Wu P, Yang Z, Busk PK, Xu Y, Yin Y. 2018. dbCAN2: a meta server for automated carbohydrate-active enzyme annotation. Nucleic Acids Res 46: W95–W101.

Zierdt CH. 1988. Blastocystis hominis, a long-misunderstood intestinal parasite. Parasitol Today Pers Ed 4: 15–17.

Zierdt CH, Williams RL. 1974. Blastocystis hominis: Axenic cultivation. Exp Parasitol 36: 233– 243.

Zimin AV, Marcais G, Puiu D, Roberts M, Salzberg SL, Yorke JA. 2013. The MaSuRCA genome assembler. Bioinformatics 29: 2669–2677.

