## Supplementary figures and images for "Contiguous and complete assemblies of *Blastocystis* gut microbiome-associated protists reveal evolutionary diversification to host ecology"

### Figure S2

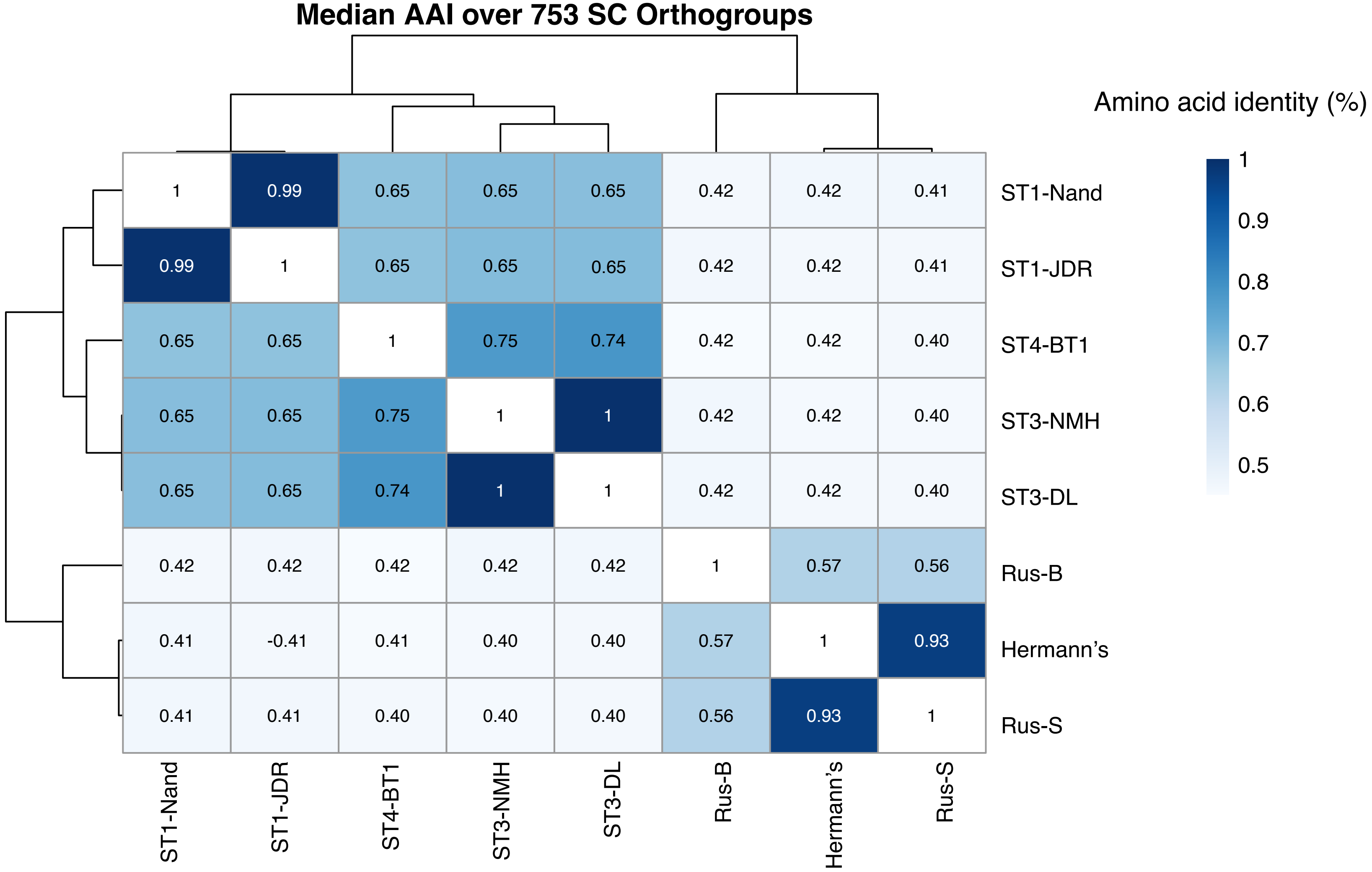

### Table S1

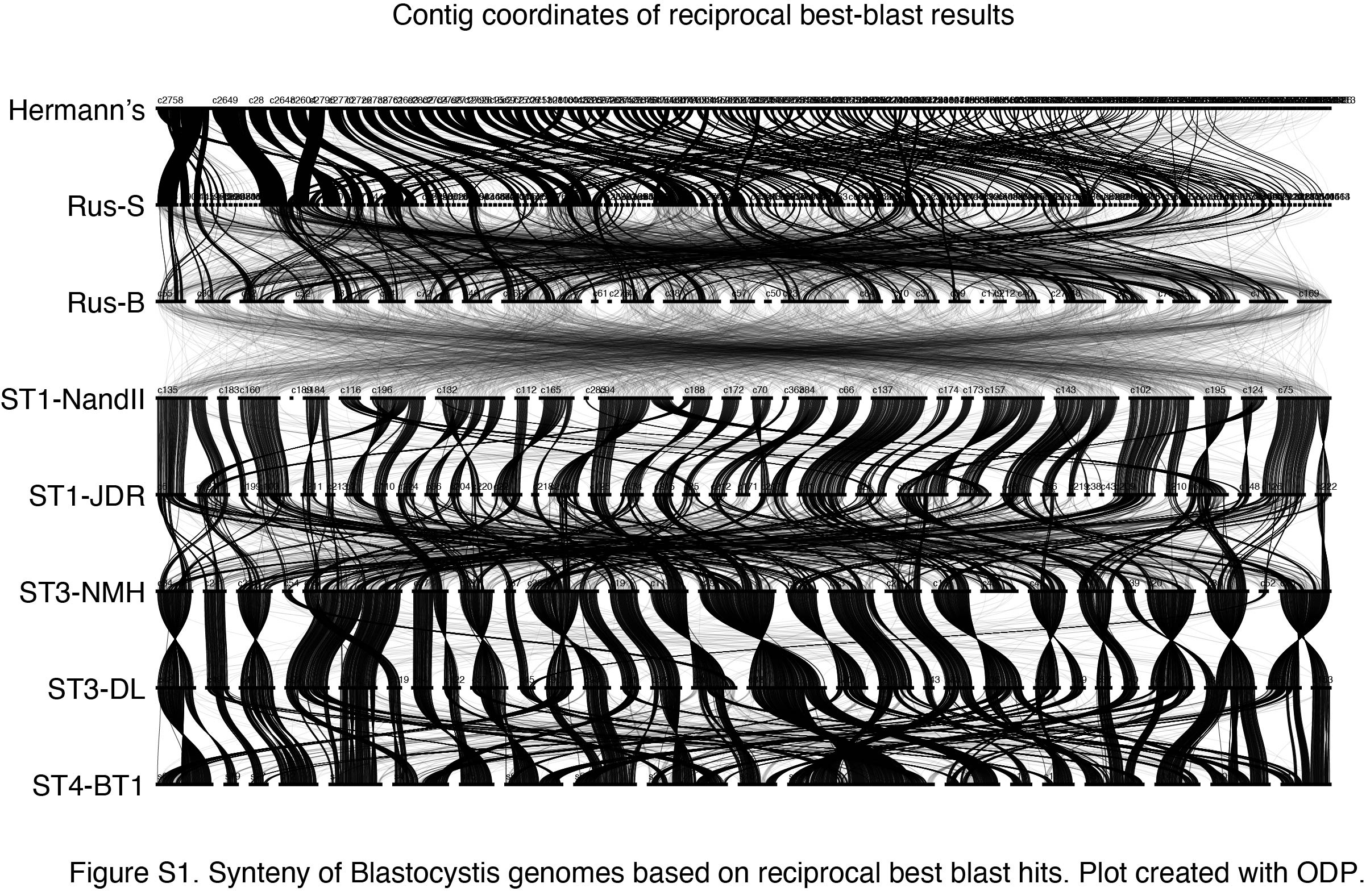
