## Supplementary material for "Contiguous and complete assemblies of *Blastocystis* gut microbiome-associated protists reveal evolutionary diversification to host ecology": Figure S3

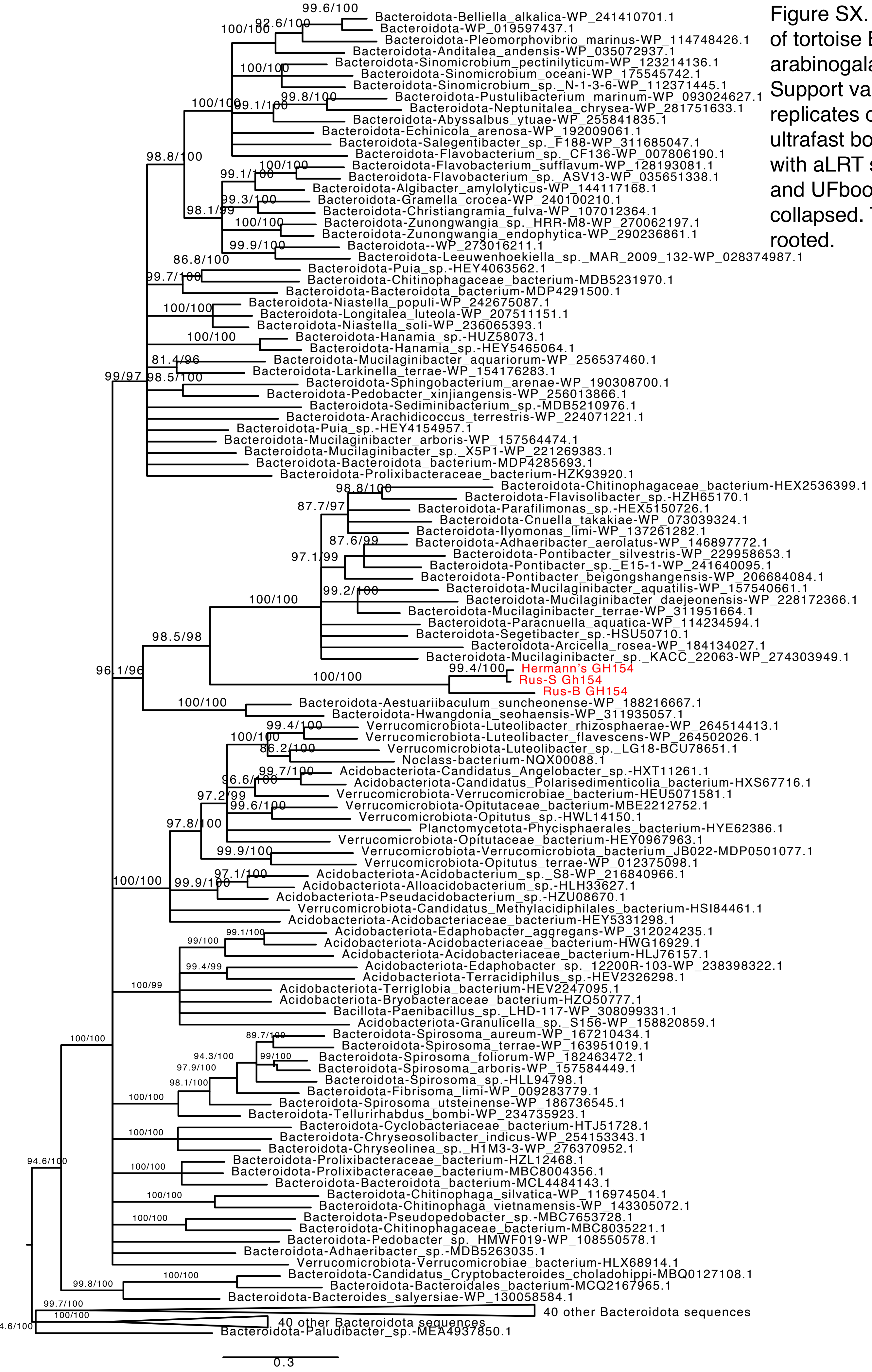


Figure SX. Gene phylogeny of tortoise Blastocystis GH154 arabinogalactanase enzyme. Support values indicate 1,000 replicates of SH-aLRT test and ultrafast bootstrap. Branches with aLRT support <80 and UFboot support <90 are collapsed. Tree is midpoint rooted.
